# SparseSeg: Target-Conditioned Discovery Segmentation of Cryo-Volume Electron Microscopy Under Sparse Annotation

**DOI:** 10.64898/2026.07.13.738355

**Authors:** Bowen Shi, Yanjun Li, Qi Ouyang, Yanan Zhu

## Abstract

Cryo-volume electron microscopy (cryo-vEM) enables near-native visualization of cellular ultrastructure, but its broad use is limited by low image contrast and the high cost of dense voxel-level annotation. Existing automated segmentation methods often generalize poorly across cell types, organelles, and imaging conditions. Here, we introduce SparseSeg, a target-conditioned, sparsity-driven segmentation framework that treats organelle segmentation as a discovery process rather than a closed-set classification task. SparseSeg uses a small number of context-specific exemplars to iteratively propagate reliable supervision through the volume. It combines sparse patch-based sampling, a multi-kernel U-Net, and geometry-consistent refinement to expand accurate segmentation while suppressing context-dependent false positives. Across serial cryo-FIB–SEM and conventional vEM datasets, SparseSeg achieves robust segmentation under extreme sparse annotation, including settings with less than 1% labeled slices. This framework reduces annotation burden while preserving morphological fidelity for quantitative cryo-vEM analysis.

## Introduction

Three-dimensional electron microscopy (3D EM) has fundamentally transformed the study of cellular organization by enabling direct visualization of organelles, membranes, and macromolecular assemblies within intact cells and tissues. In particular, volume electron microscopy (vEM), including focused ion beam scanning electron microscopy (FIB–SEM), serial block-face SEM (SBF–SEM), and related approaches, provides nanoscale reconstructions across entire cells and extended tissue volumes, revealing spatial relationships that are inaccessible to conventional two-dimensional imaging [1–5].

Recent advances in vEM have enabled comprehensive, whole-cell organelle atlases and large-scale reconstructions that support quantitative analysis of cellular architecture at unprecedented scale [6, 7]. These efforts demonstrate the transformative potential of automated segmentation for integrative structural cell biology. At the same time, they expose a fundamental limitation: generating dense voxel-level annotations for volumetric EM data remains extraordinarily labor-intensive, often requiring months of expert manual effort for a single cell [6, 8]. This annotation burden constrains scalability, limits reproducibility, and places quantitative vEM analysis beyond the reach of many biological studies.

Cryo-volume electron microscopy (cryo-vEM)—in particular serial cryo-FIB–SEM—offers a unique opportunity to overcome many limitations of conventional vEM by imaging biological specimens in a near-native, vitrified state [9–11]. By combining high-pressure freezing with cryogenic FIB milling and SEM imaging, cryo-vEM preserves native membrane topology, organelle morphology, and macromolecular organization without distortions introduced by chemical fixation, dehydration, or heavy-metal staining, which is critical for studying subtle ultrastructural features [9, 12].

Despite its biological importance, cryo-vEM presents a severe analytical challenge. Modern cryo-FIB–SEM experiments routinely generate terabyte-scale datasets comprising hundreds to thousands of serial sections [13]. Manual segmentation of such data is prohibitively time-consuming and difficult to reproduce, rendering systematic and statistically powered biological analyses impractical. Consequently, much of the rich structural information captured by cryo-vEM remains underutilized.

Deep learning–based segmentation has emerged as a powerful approach for automated EM analysis and has achieved impressive performance on conventionally prepared, heavily stained datasets [6, 14–16]. However, segmentation models trained on conventional vEM often fail when applied to cryo-vEM data. Cryogenic preserved images are characterized by low contrast and reduced signal-to-noise ratios, leading to unreliable segmentation even for relatively well-defined organelles such as mitochondria [8, 17]. We argue that this limitation reflects not a failure of deep learning per se, but a fundamental mismatch between the contrast mechanisms of cryo-native imaging and the assumptions encoded in models trained on conventional vEM data.

More fundamentally, the appearance and segmentability of a given organelle are strongly context-dependent. Even within the same organelle class, the contrast, topology, and surrounding ultrastructure vary substantially across cell types, physiological states, and imaging conditions. Annotation workflows—including semi-automated approaches such as ilastik [18]—can further introduce variability through user-dependent feature selection and labeling strategies. As a result, segmentation models trained to exhaustively recognize predefined organelle classes under one context often generalize poorly to new datasets, even when organelle morphology is conserved.

These observations motivate a shift in how organelle segmentation is formulated for cryo-vEM. Rather than treating segmentation only as a closed-set classification problem requiring dense and exhaustive annotations, we propose framing it as a *target-conditioned discovery process*. In this paradigm, a small number of context-specific exemplars are used to condition the model to identify and propagate additional instances of the same structure within a given dataset. This formulation is not intended to replace generalist segmentation models, foundation models, or domain adaptation strategies. Instead, it provides a complementary mechanism for rapid dataset-specific adaptation when a new imaging condition, cell type, or target structure is encountered. By enabling users to generate reliable target-specific supervision with minimal annotation effort, SparseSeg can also support downstream fine-tuning or adaptation of more general segmentation models.

Here, we introduce SparseSeg, a target-conditioned, sparsity-driven segmentation framework designed explicitly for cryo-vEM. Our approach integrates sparse patch-based sampling, a multi-kernel U-Net architecture optimized for cryo-native contrast, and iterative geometry-consistent refinement to progressively expand reliable supervision from minimal annotations. We demonstrate robust segmentation performance under extreme annotation scarcity across multiple cell types, organelles, and imaging conditions. By enabling accurate and reproducible segmentation with minimal manual input, this framework establishes a practical and scalable foundation for quantitative cryo-vEM analysis and broadens access to *in situ* ultrastructural biology.

## Results

### Framework overview and workflow

SparseSeg is designed to recover biological targets from extremely sparse user annotations. Rather than relying on dense voxel-level supervision or pre-trained models, the framework uses sparse annotations as conditioning signals to guide representation learning, structural propagation, and iterative target discovery.

The overall workflow comprises three tightly integrated modules: (i) target-conditioned sparse supervision, (ii) context- and geometry-aware representation learning, and (iii) iterative discovery and refinement. An overview of the complete pipeline is shown in Fig. 1.

**Figure 1.**
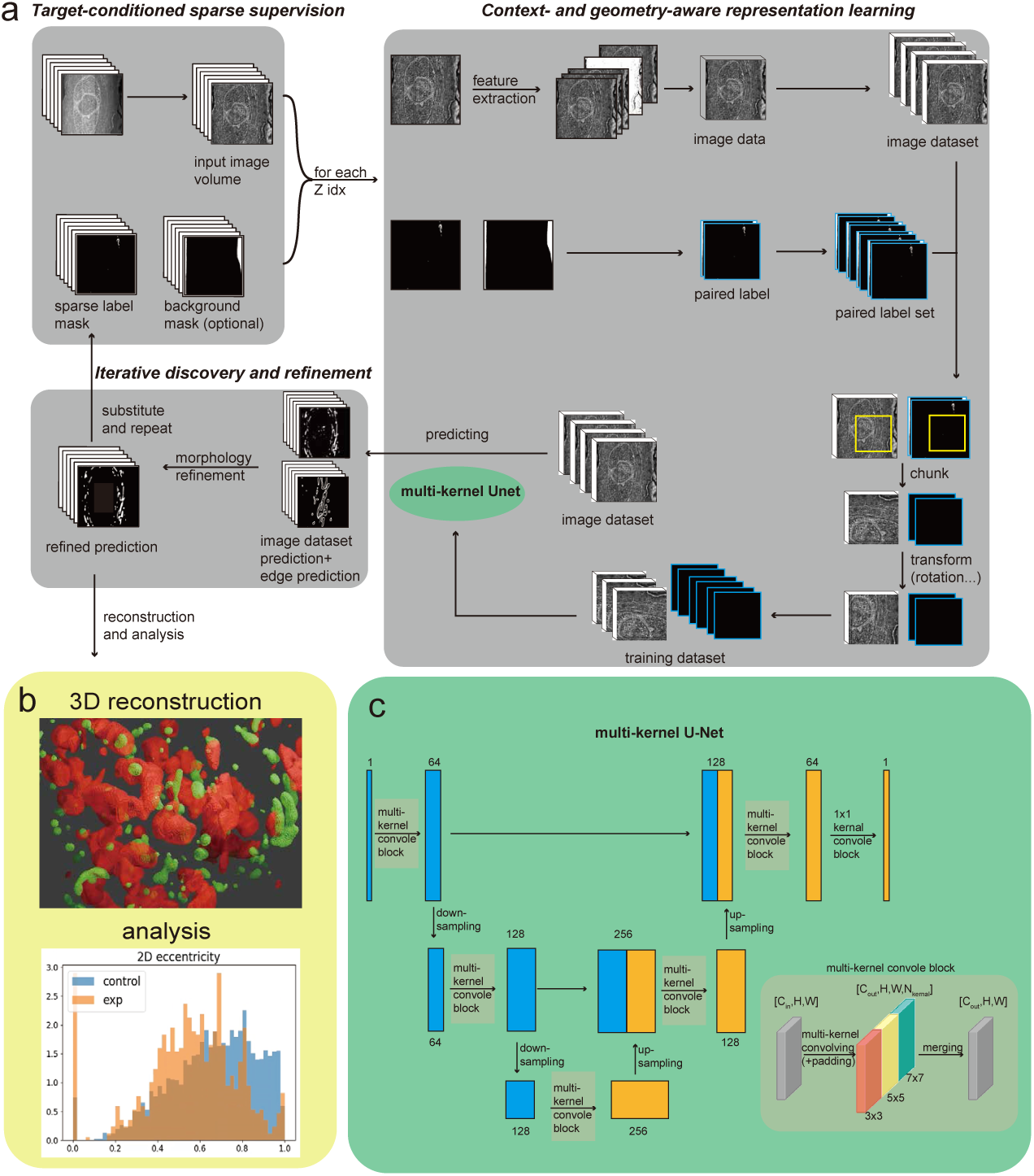
Overview of the proposed segmentation framework and model architecture. **a,** Schematic of the overall workflow. The framework consists of three core modules: target-conditioned sparse supervision, context- and geometry-aware representation learning, and iterative discovery and refinement (highlighted in gray). **b,** Example downstream applications enabled by the framework, including three-dimensional reconstruction and quantitative morphometric analysis of segmented organelles. **c,** Architecture of multi-kernel U-Net, which integrates parallel convolutional branches with multiple receptive fields to capture fine membrane details, intermediate organelle morphology, and broader contextual structures in cryo-volume electron microscopy data.

The first module uses target-conditioned sparse supervision.

Raw vEM datasets often exhibit substantial variability in image contrast, spatial scale, and inter-slice alignment due to differences in microscope settings, acquisition conditions, and beam-induced drift. To standardize the inputs, an optional preprocessing step is applied, as described in the Methods.

Sparse user annotations form the core supervision signal of SparseSeg. Users are required to annotate only a small number of representative ROIs, typically fewer than 10 two-dimensional regions, indicating partial locations of the target organelle in the raw image volume *V*. These sparse positive annotations are encoded as a binary sparse label mask *M*. Optionally, users may provide background negative ROIs marking regions where the target structure is confidently absent; these annotations are encoded as a binary negative annotation mask *N*. Unless otherwise specified, *V*, *M*, and *N* denote the input image volume, the sparse positive label mask, and the optional negative annotation mask throughout the manuscript.

This target-conditioned supervision enables the model to adapt to dataset-specific context and imaging conditions with minimal labeling effort.

The second module uses context- and geometry-aware representation learning.

For a single three-dimensional image volume *V* ∈ ℝ*^D×H×W^*, SparseSeg constructs a corresponding feature volume *F* ∈ ℝ*^D×C×H×W^*, where *D*, *H*, *W*, and *C* denote the image depth, height, width, and feature dimension, respectively. Although the sparse annotation masks *M* and *N* are stored as volumetric stacks with the same spatial dimensions as *V*, the supervision is provided at the two-dimensional ROI level. In other words, users annotate sparse two-dimensional regions on selected *z*-slices, and these annotations are stored in the corresponding slices of *M* and *N*. The unannotated slices and pixels are treated as unlabeled rather than dense negative supervision.

Given the feature volume *F* and the sparse supervision masks (*M, N*), training samples are generated by sampling two-dimensional spatial patches from selected slices. Patch centers are anchored on annotated positive regions, with a small fraction of patches sampled from user-provided negative regions when available (default 3%). Each training sample therefore consists of a feature patch *F* [*z*] from one slice, together with the corresponding two-dimensional supervision masks *M* [*z*] and *N* [*z*]. Local axial context can be incorporated into *F* [*z*] during feature construction, but the manual supervision itself remains sparse and two-dimensional.

The positive, negative, and soft-negative masks are cropped accordingly, and an additional edge supervision channel is derived from the two-dimensional positive mask using a one-pixel-wide morphological boundary. This edge channel explicitly emphasizes thin membranes and weakly contrasted boundaries characteristic of cryo-vEM data. We further generate and crop coarse edge and area reference maps for use in the consistency losses, as described in the Methods and Supplementary.

Prediction and training are performed using the multi-kernel U-Net architecture described in Fig. 1c. The network is applied slice-wise to feature patches, and the resulting predictions from all slices are assembled into a three-dimensional segmentation volume. Each convolutional block contains parallel branches with different kernel sizes, enabling simultaneous encoding of fine membrane boundaries, intermediate organelle morphology, and larger contextual structures.

Representation learning is guided by a composite objective that integrates masked classification loss, region consistency and contrast constraints, image-guided smoothness, and sparsity regularization as shown in Supplementary.

The third module performs iterative discovery and refinement.

Although the trained multi-kernel U-Net can already produce dense slice-wise probability maps that are assembled into a volumetric prediction, raw predictions may contain spurious regions or fragmented structures, particularly in low-contrast cryo-vEM data. To address this, a geometry-consistent refinement stage is introduced to enable robust target discovery and iterative self-improvement.

Predicted masks can be decomposed into region interiors and boundary-like components. Region interiors are extracted and characterized using geometric descriptors and moment invariants. Each predicted region interior is compared with reference regions extracted from the sparse annotation mask *M*, and only components with sufficiently similar shape statistics are retained. In parallel, predicted boundary-like components are used to validate region interiors via bounding-box overlap constraints, enforcing spatial consistency between region interiors and boundary-like components.

Refined predictions that satisfy both shape similarity and spatial consistency criteria are promoted to reliable labels and merged with existing annotations. These augmented labels are then fed back into the training pipeline for subsequent iterations. Through this iterative discovery-and-refinement process, the model progressively expands supervision coverage, improves segmentation quality, and remains robust to error accumulation.

Together, these three modules enable effective propagation of sparse annotations into complete, geometry-consistent organelle segmentations, even under severe annotation scarcity, substantial domain shift, and varying contextual conditions.

### Few-shot segmentation under extremely sparse supervision

To evaluate the generalizability of our framework in a few-shot regime, we applied it to multiple independent vEM datasets spanning diverse organisms, imaging modalities, and biological targets. This experiment assesses the ability of the framework to adapt to different datasets using only a small number of annotations (typically fewer than 10 annotated ROIs per dataset).

We first evaluated few-shot generalization across diverse biological targets and vEM modalities.

We first focused on mitochondrial segmentation, as mitochondria are among the most widely studied yet morphologically diverse organelles in EM analysis. We evaluated performance on several public EMPIAR datasets acquired using both conventional and cryogenic vEM modalities, including an interphase HeLa cell (Fig. 2a) imaged by FIB–SEM (EMPIAR-10311) [19], plant root tissue (Fig. 2b) imaged by serial block-face SEM (EMPIAR-10442) [20], and cryo serial FIB–SEM datasets of mouse brain tissue (Fig. 2c), *Saccharomyces cerevisiae* (Fig. 2d), Vero cells (Fig. 2e), and HeLa cells (Fig. 2f) (EMPIAR-11415, EMPIAR-11416, EMPIAR-11417, and EMPIAR-11419) [21].

**Figure 2.**
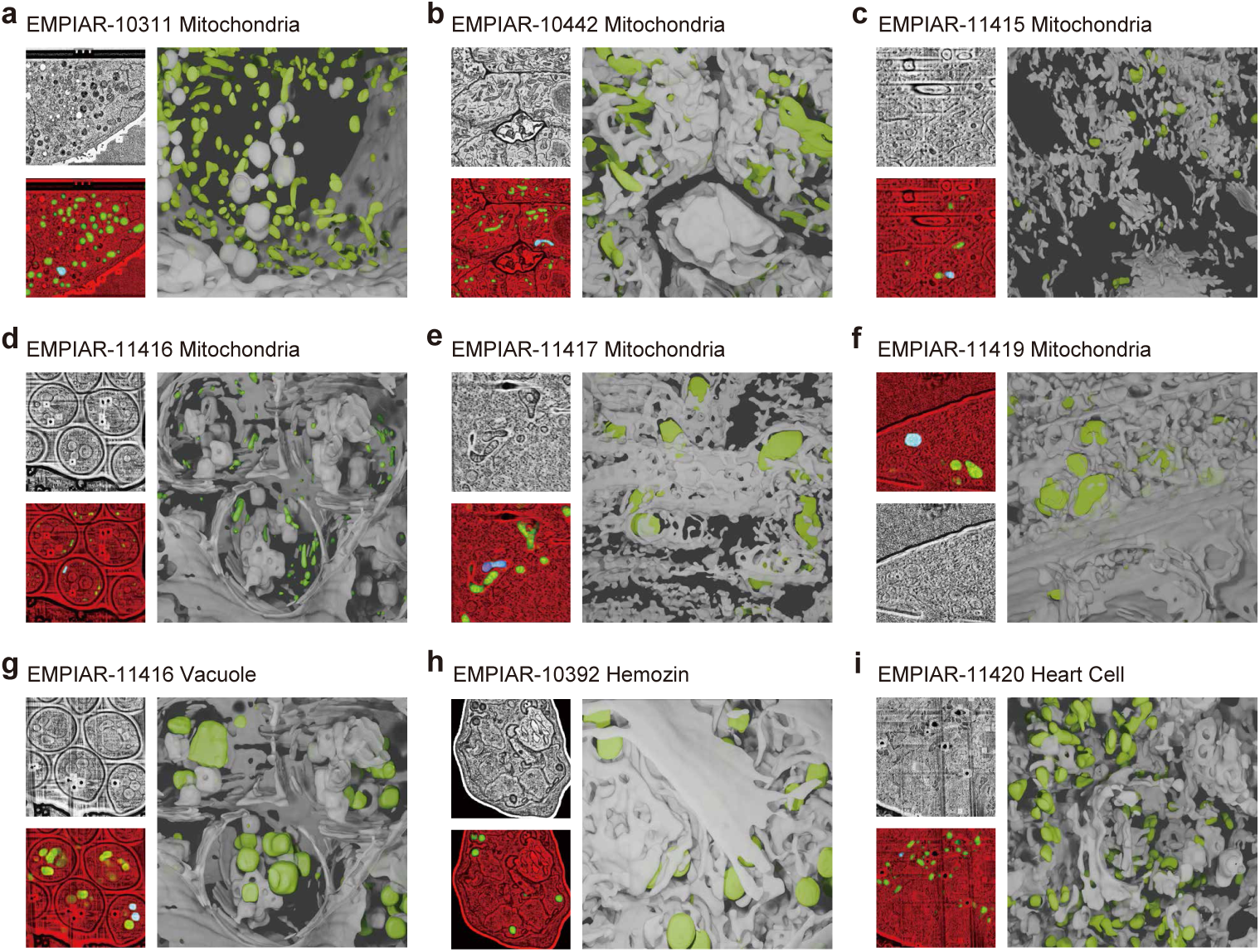
Few-shot identification of diverse organelles and cell types across heterogeneous volume electron microscopy datasets. Representative two-dimensional slices show raw images, model predictions, and corresponding three-dimensional reconstructions. In the prediction overlays, red denotes background, green denotes predicted target structures, and blue denotes sparse annotation when present. For three-dimensional visualization, the grayscale volume shown in white is generated by isosurface rendering of the raw image intensity to provide anatomical context only, whereas the green structures correspond to model-predicted targets. **a,** Mitochondrial segmentation in an interphase HeLa cell imaged by focused ion beam scanning electron microscopy (FIB–SEM; EMPIAR-10311). **b,** Mitochondrial segmentation in *Arabidopsis thaliana* root tissue imaged by serial block-face scanning electron microscopy (SBF–SEM; EMPIAR-10442). **c,** Mitochondrial segmentation in mouse brain tissue acquired by cryo serial FIB–SEM (EMPIAR-11415). **d,** Mitochondrial segmentation in *Saccharomyces cerevisiae* using cryo serial FIB–SEM (EMPIAR-11416). **e,** Mitochondrial segmentation in Vero cells imaged by cryo serial FIB–SEM (EMPIAR-11417). **f,** Mitochondrial segmentation in HeLa cells imaged by cryo serial FIB–SEM (EMPIAR-11419). **g,** Vacuole segmentation in *Saccharomyces cerevisiae* (EMPIAR-11416). **h,** Hemozoin crystal segmentation in *Plasmodium falciparum* across cytokinesis (EMPIAR-10392). **i,** Cardiomyocyte segmentation in mouse heart tissue (EMPIAR-11420). Together, these results demonstrate robust generalization across organisms, imaging modalities, and biological targets under extreme annotation sparsity.

For each dataset, supervision was provided by manually annotating fewer than 10 two-dimensional mitochondrial regions across the entire volume (highlighted in blue in Fig. 2). These annotations were sparse, non-contiguous, and confined to a small subset of slices within volumes typically comprising approximately 50 to more than 1500 serial sections. Consequently, the fraction of annotated slices is extremely small relative to the total volume, reaching fewer than 0.1% in the *Saccharomyces cerevisiae* cryo serial FIB–SEM dataset (EMPIAR-11416). Such annotations can be completed within 1–2 minutes using the proposed pipeline (see Methods). Despite this minimal supervision, the model successfully recovered the majority of mitochondrial structures throughout each volume, as indicated by the predicted regions shown in green in Fig. 2a–f.

We further extended this analysis beyond mitochondria to encompass diverse biological targets and vEM imaging modalities. As shown in Fig. 2g, the model successfully segments vacuoles in *Saccharomyces cerevisiae* (EMPIAR-11416) [21]. Figure 2h shows segmentation of membrane-free hemozoin crystals in the malaria parasite *Plasmodium falciparum* (EMPIAR-10392) [22], a dataset acquired using conventional FIB–SEM with heavy-metal staining. These structures exhibit ultrastructural and contrast characteristics that differ markedly from mitochondria. Accurate identification of such targets demonstrates that the framework is not restricted to specific organelles, but operates robustly across distinct structural classes, including both membrane-enclosed and membrane-free features, in cryo and conventional vEM datasets.

Furthermore, Fig. 2i illustrates application of the same framework to cell identification in cryo serial FIB–SEM data of mouse heart tissue (EMPIAR-11420) [21], where spatial scale differs substantially from the previous datasets. Despite these differences, sparsely labeled cardiomyocytes were reliably identified and delineated across a multicellular tissue volume.

Taken together, these results demonstrate that the model generalizes across vEM modalities, including cryogenic FIB–SEM, conventional FIB–SEM, and SBF–SEM, as well as across diverse biological targets ranging from subcellular organelles to cell types. By conditioning segmentation on sparse, dataset-specific annotations rather than modality-specific appearance priors, the framework provides a flexible target-conditioned strategy for vEM analysis that can be rapidly adapted to different imaging conditions using sparse dataset-specific annotations.

We next assessed three-dimensional reconstruction of diverse organelle structures across cell types.

To further assess robustness and structural fidelity, we reconstructed whole three-dimensional cells containing multiple organelles across different vEM datasets from the Janelia COSEM project, including Jurkat T cells, HeLa cells, and macrophages. All datasets were prepared by high-pressure freezing, followed by freeze substitution with heavy-metal staining and resin embedding, and subsequently imaged using FIB–SEM.

Figure 3 presents representative two-dimensional slices together with corresponding three-dimensional renderings of multiple organelles, including mitochondria, nucleus, Golgi apparatus, lysosomes, endosomes, and endoplasmic reticulum (ER). These organelles collectively exhibit diverse ultrastructural morphologies, including elongated and branched tubular networks (mitochondria), thin sheet-like and reticular membranes (ER), compact stacked cisternae (Golgi apparatus), large enclosed volumes (nucleus), and small, heterogeneous vesicular structures (lysosomes and endosomes).

**Figure 3.**
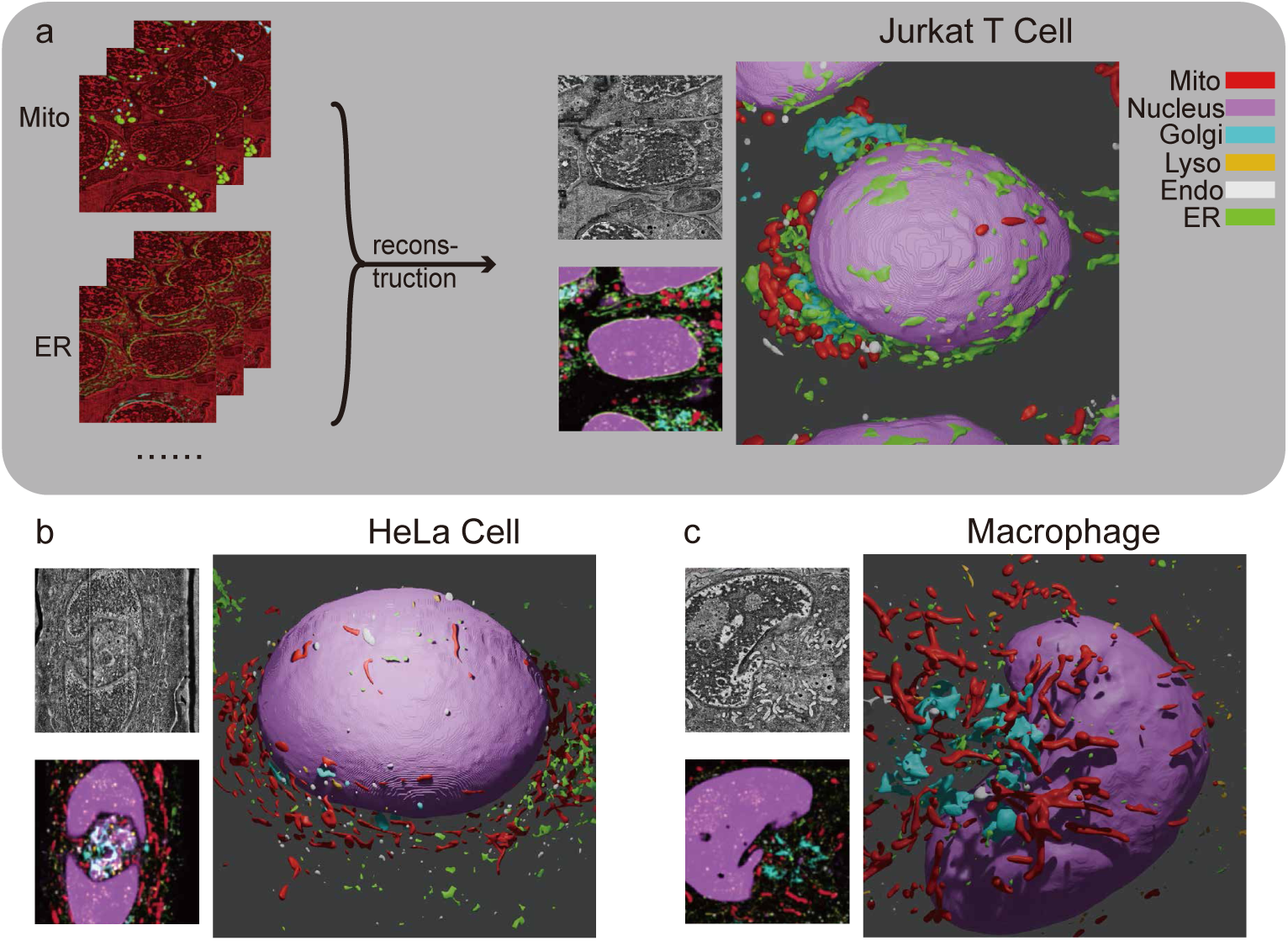
Three-dimensional reconstruction of diverse organelle structures from model-predicted segmentations. **a,** Representative two-dimensional slices used for volumetric reconstruction, together with the corresponding bright-field reference images, composite slice overlays, and three-dimensional renderings from a Jurkat T cell. Multiple organelles are accurately reconstructed, including mitochondria, endoplasmic reticulum, Golgi apparatus, lysosomes, nucleus, and endosomes. The reconstructed geometries preserve organelle continuity, topology, and native spatial organization, enabling direct visualization and quantitative analysis of cryo-volume electron microscopy ultrastructure. **b,** Three-dimensional reconstructions from a HeLa cell volume, together with corresponding bright-field references and composite slice overlays, highlighting consistent volumetric reconstruction quality and generalization across distinct cell types. **c,** Three-dimensional organelle reconstructions from a macrophage dataset, shown alongside corresponding bright-field images and composite slice views, demonstrating robust segmentation and volumetric reconstruction in a morphologically complex mammalian cell.

In Jurkat T (Fig. 3a) and HeLa cells (Fig. 3b), the model accurately captures elongated and branched mitochondria, making their interconnected network clearly visible. The ER is reconstructed as both tubular networks and sheet-like cisternal structures, which are particularly challenging due to their thin membranes and heterogeneous morphology. Golgi apparatuses are resolved as compact perinuclear stacked cisternae, clearly distinguishable from surrounding vacuoles, while lysosomes and endo-somes—characterized by small volumes and high morphological variability—are also reliably segmented.

Comparable reconstruction quality is observed in the macrophage dataset (Fig. 3c), demonstrating that the framework generalizes across cell types with substantially different cellular architectures and modes of organelle organization.

We then examined challenging cases and robustness.

The visualizations highlight several challenging segmentation scenarios, including closely apposed or touching organelles, densely packed perinuclear regions, and faint or partially discontinuous membranes typical of cryo-preserved imaging. Despite these challenges, the model successfully separates adjacent organelles and preserves their three-dimensional continuity, enabling coherent volumetric reconstructions.

Notably, the reconstructed three-dimensional renderings exhibit anatomically consistent spatial organization, such as mitochondria distributed around the nucleus and ER forming extended networks throughout the cytoplasm. These observations indicate that the model learns structurally and topologically meaningful representations, rather than producing purely voxel-wise predictions.

### Benchmarking performance

To quantitatively assess segmentation performance, we conducted benchmarking experiments on three mammalian whole-cell datasets from the Janelia COSEM project, including HeLa cells, Jurkat T cells, and macrophages, all of which provide high-quality expert-annotated ground truth.

We evaluated SparseSeg under two complementary annotation-sparsity settings. First, we used a controlled masking protocol in which a specified fraction of annotated image slices was randomly withheld to simulate incomplete supervision. This masking procedure was applied consistently across the evaluated methods. Segmentation performance was quantified using the Intersection-over-Union (IoU) metric, with precision and recall computed relative to expert manual annotations provided by the COSEM Project Team.

As shown in Fig. 4, SparseSeg consistently outperformed StarDist across all masking ratios and cell types. In the HeLa dataset, SparseSeg maintained IoU values of approximately 0.5 across all masking conditions, whereas StarDist achieved only around 0.2. SparseSeg also exhibited more stable performance across varying masking ratios, indicating enhanced robustness to incomplete supervision.

**Figure 4.**
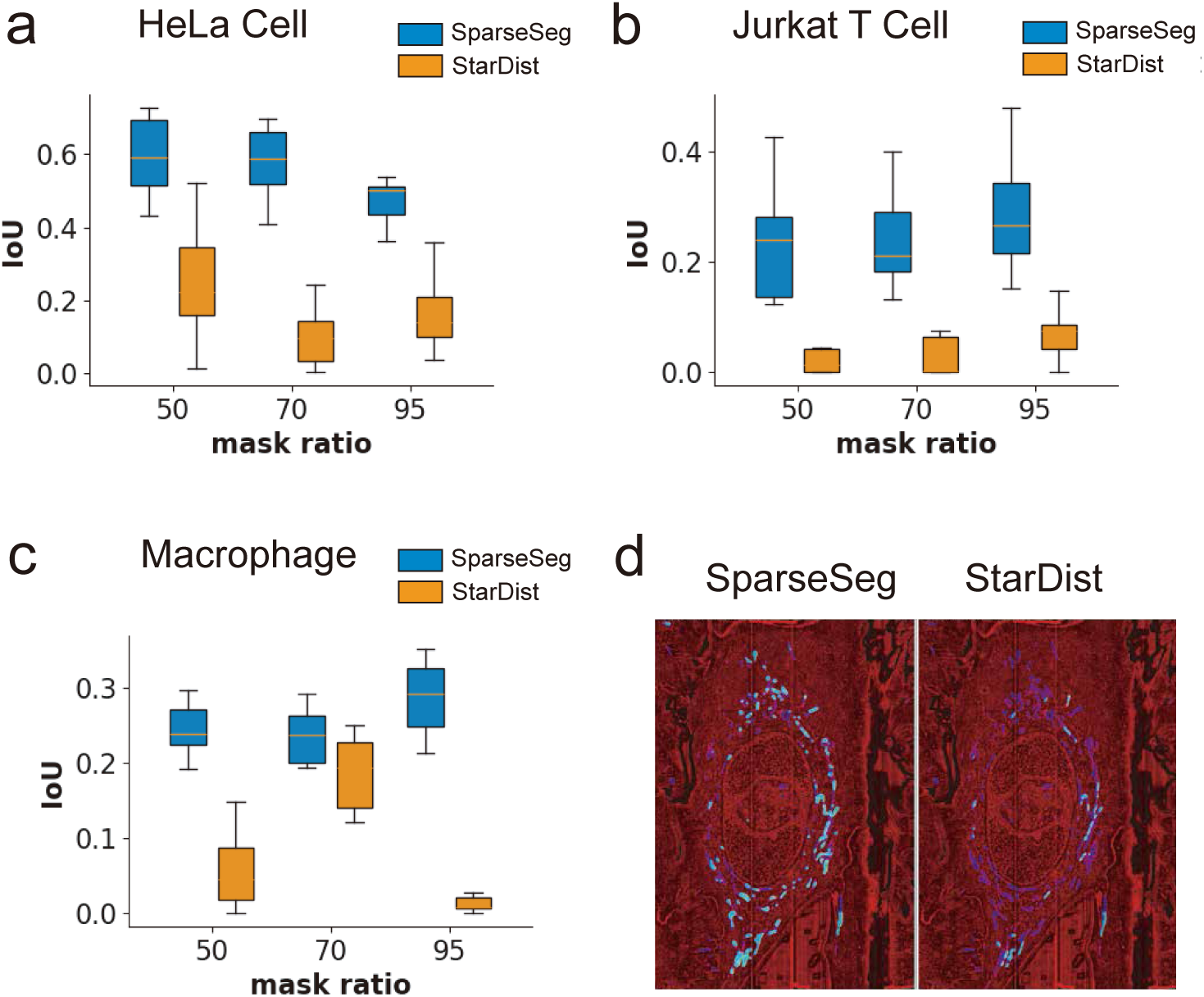
Quantitative benchmarking of mitochondrial segmentation performance under controlled label sparsity. **a,** Segmentation performance comparison for 10 trials on the HeLa cell dataset as a function of masking ratio, defined as the fraction of ground-truth slices withheld during training. **b,** Corresponding results on the Jurkat T cell dataset. **c,** Corresponding results on the macrophage cell dataset. **d,** Representative segmentation results at a masking ratio of 95% for SparseSeg and StarDist. Cyan regions indicate true positives, blue regions denote false negatives, corresponding to ground-truth regions not recovered by the model, and green regions correspond to false positives.

Second, we further evaluated SparseSeg under a more challenging ROI-level sparse-annotation setting, where only 1, 5, or 10 ROIs were randomly selected as positive target labels. We compared SparseSeg with a broad range of benchmark models under the same annotation setting, including MitoNet [23], nnU-Net [24], COSEM-style 2D/3D U-Net baselines for cellular EM segmentation [6], DeePiCt [25], StarDist [26, 27], and SparseSeg-ViT based on the Vision Transformer architecture [28]. Because most existing segmentation models are not specifically designed for such sparse-label training, their absolute IoU values can be relatively low in this regime. Therefore, we used normalized relative IoU as the main metric to facilitate comparison across models and sparsity levels.

As shown in Fig. 5a, SparseSeg and SparseSeg-ViT achieved the best performance across all tested annotation levels. The dashed line in Fig. 5a indicates the normalized relative IoU obtained using the pretrained MitoNet model without task-specific sparse-label fine-tuning. This result suggests that the target-conditioned sparse segmentation framework is robust under extremely limited supervision. The strong performance of SparseSeg-ViT further indicates that the overall SparseSeg framework can be extended to different backbone architectures, rather than being restricted to the original multi-kernel U-Net implementation.

**Figure 5.**
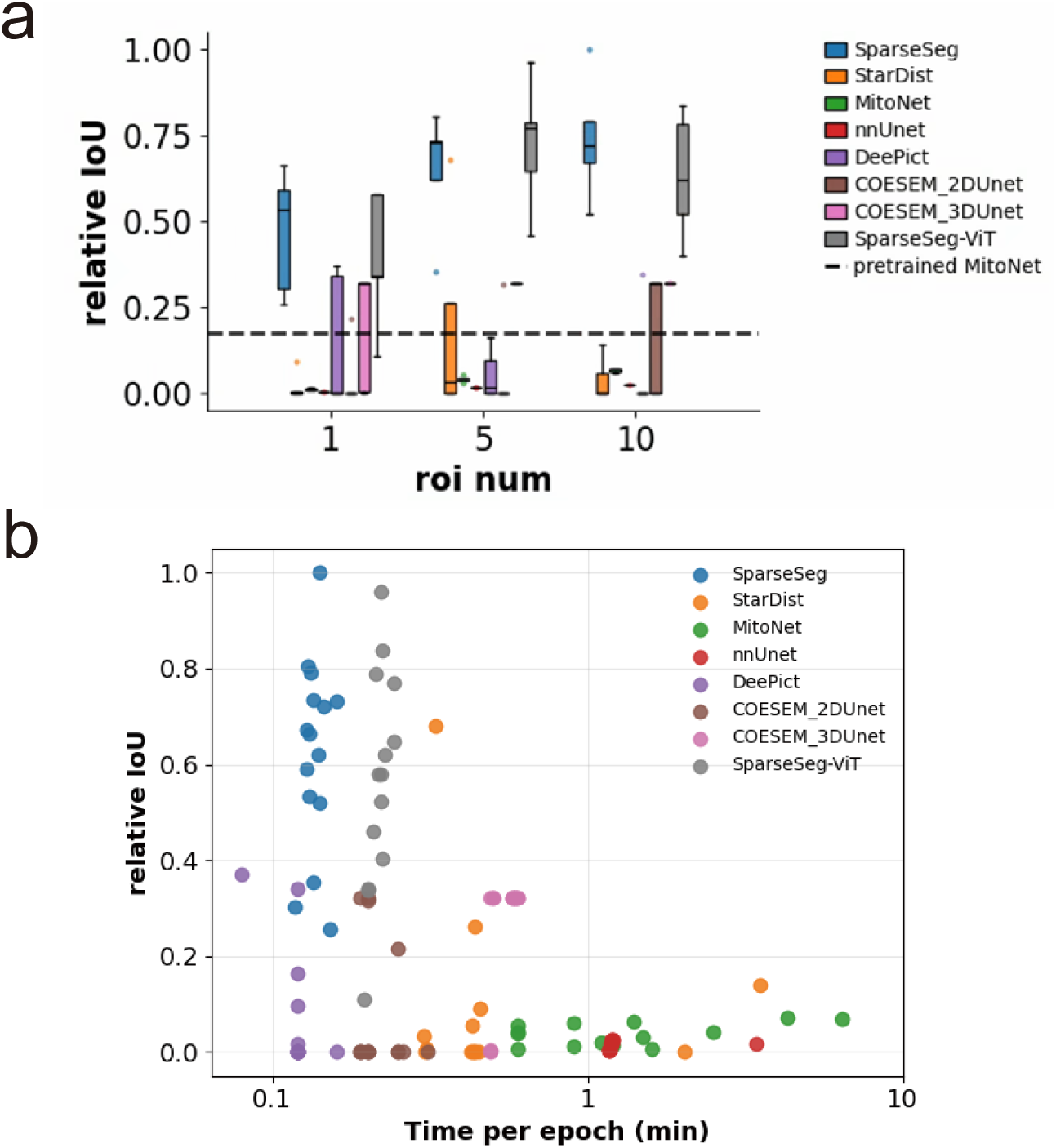
Quantitative benchmarking of mitochondrial segmentation under region-of-interest-level sparse annotations. **a,** Normalized relative intersection over union (IoU) across different models using 1, 5, or 10 randomly selected regions of interest (ROIs) as positive target labels. The dashed line indicates the normalized relative IoU obtained using the pretrained MitoNet model without task-specific sparse-label fine-tuning. **b,** Training time per epoch for different benchmark models measured on the same workstation.

We also measured the training time per epoch for each model on the same workstation. As shown in Fig. 5b, DeePiCt and SparseSeg required the shortest training time per epoch, followed by SparseSeg-ViT and COSEM-2D U-Net. These results indicate that SparseSeg provides competitive segmentation performance under sparse annotations while maintaining relatively low computational cost.

Across both benchmarking settings, SparseSeg achieved strong segmentation performance under limited supervision. These results indicate that the advantages of SparseSeg are not limited to a specific sparsification protocol, but extend from controlled masking of dense annotations to extreme ROI-level sparse supervision.

We next performed a quantitative comparison with existing segmentation methods. We compared our model (SparseSeg) against a widely used segmentation approach originally developed for conventional EM data: StarDist [29], a shape-based instance segmentation method.

Mitochondria were selected as the initial benchmarking target due to their complex and highly variable morphology. Expert manual segmentations provided by the COSEM Project Team were used as the gold-standard reference for quantitative evaluation. Details of training and test data construction, including masking procedures, are provided in Supplementary.

As shown in Fig. 4 and Supplementary Fig.1, SparseSeg consistently outperformed StarDist across all masking ratios and cell types. In the HeLa dataset, SparseSeg maintained IoU values of approximately 0.5 across all masking conditions, whereas StarDist achieved only around 0.2. Moreover, SparseSeg exhibited substantially more stable performance across varying masking ratios compared to StarDist, as evidenced by results across multiple trials (Fig. 4), indicating enhanced robustness to annotation sparsity.

Across additional benchmarks, SparseSeg consistently outperformed StarDist in the majority of cases, demonstrating superior robustness to incomplete observation, and morphological variability. Collectively, these results indicate that the advantages of SparseSeg are not limited to mitochondria or a single cell type, but generalize across diverse cellular contexts and organelle classes (Fig. 4 and Supplementary Fig.1).

We also tested context-dependent generalization to evaluate the need for dataset-specific adaptation.

Figure 6 evaluates model performance across different cellular contexts and annotation sparsity levels. Models were trained to segment mitochondria using data from a single cell type and subsequently tested on unseen datasets from other cell types, including HeLa cells, Jurkat T cells, and macrophage, under varying degrees of label sparsity.

**Figure 6.**
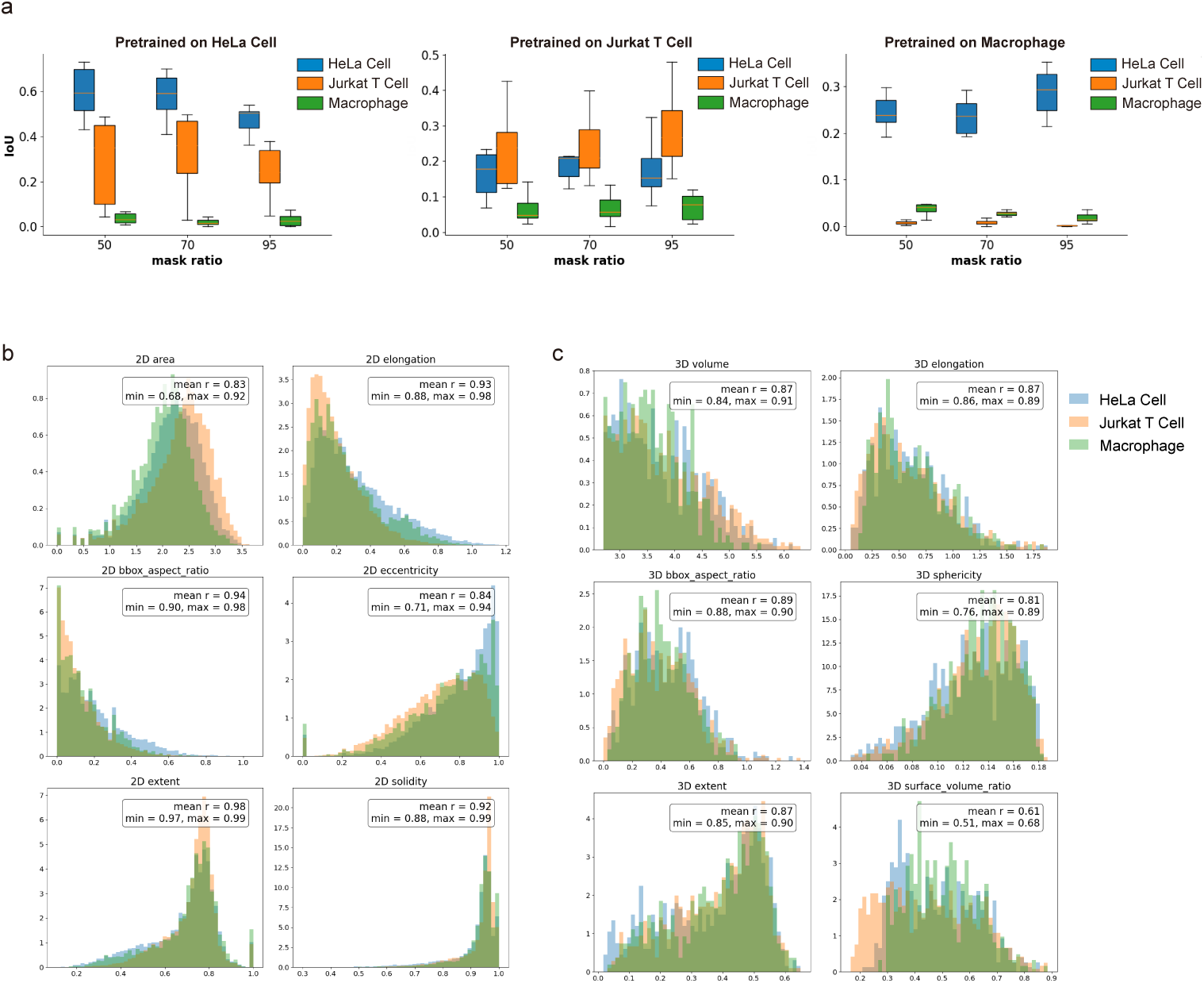
Generalization of mitochondrial segmentation across datasets, cell types, and annotation sparsity levels. **a,** Cross-dataset generalization performance of SparseSeg models pretrained on different cell types, including HeLa cells, Jurkat T cells, and macrophages, under varying degrees of label sparsity. Each model is pretrained using sparsely annotated mitochondria from one dataset and evaluated on unseen datasets. **b,** Two-dimensional mitochondrial morphological statistics across datasets, including shape descriptors such as eccentricity and aspect ratio. **c,** Three-dimensional mitochondrial morphological statistics derived from volumetric reconstructions. Despite substantial differences in segmentation generalization, mitochondrial morphology remains highly conserved across datasets, indicating that generalization failure is driven primarily by differences in cellular context and imaging characteristics rather than intrinsic organelle shape.

Despite being trained on the same organelle class, segmentation performance is not uniformly preserved across datasets (Fig. 6a). This degradation cannot be explained solely by differences in mitochondrial morphology. Quantitative analysis shows that key shape descriptors, such as eccentricity, remain highly consistent across datasets, with minimal pairwise correlation coefficients exceeding 0.7. Instead, the observed performance gap is primarily driven by differences in the surrounding cellular context.

Specifically, models trained on HeLa and Jurkat T cells datasets exhibit strong cross-generalization, maintaining comparable prediction accuracy when applied to each other’s data. In contrast, models trained on macrophage datasets fail to reliably identify mitochondria in HeLa or Jurkat T cells. This asymmetry reflects substantial differences in cellular organization and ultrastructural context between macrophages and the other cell types, despite the conserved intrinsic morphology of mitochondria.

These results provide empirical support for our central design principle: the appearance and segmentability of the same organelle are strongly context-dependent, varying with cell type and imaging conditions. Consequently, training a model to recognize a specific organelle from densely annotated data in one context does not necessarily guarantee robust performance across datasets. This observation does not exclude the possibility that larger and more diverse training datasets, domain adaptation, or foundation segmentation models could improve cross-domain generalization. Rather, our results suggest that sparse target-conditioned adaptation provides a practical and annotation-efficient complementary strategy, particularly when broadly generalizable models are not available or when new dataset-specific structures need to be analyzed. By leveraging a small number of sparsely labeled target structures, SparseSeg can rapidly adapt to local imaging context and identify additional instances within the same dataset. This target-driven, context-adaptive strategy helps accommodate variability introduced by biological differences, imaging conditions, and annotation workflows, including semi-automated approaches such as ilastik, which can further influence labeling characteristics.

Together, these findings highlight the importance of sparse, adaptive supervision for achieving robust generalization in vEM segmentation and underscore the limitations of conventional fully supervised approaches in highly heterogeneous imaging settings.

### High-throughput quantification reveals native-state biology

To demonstrate that SparseSeg supports downstream analysis beyond visual inspection, we applied it to quantitative mitochondrial analysis in a cryo-FIB–SEM dataset from Leigh syndrome patient cells [12]. Using mitochondrial segmentations generated by SparseSeg (Fig. 7a,b), we performed three-dimensional and two-dimensional morphometric analyses to compare mitochondrial organization between patient-derived and control examples. Because matched multi-cell datasets with control and Leigh syndrome annotations are unavailable, this analysis was performed at the segmented-object level rather than at the biological-replicate level. Accordingly, these results should be interpreted as an illustrative object-level application of SparseSeg, rather than as a definitive biological-replicate-level or population-level comparison. For statistical analysis, each reconstructed 3D mitochondrial object or segmented 2D mitochondrial cross-section was treated as one measurement. The 3D analysis included *n* = 176 control mitochondrial objects and *n* = 69 patient-derived mitochondrial objects, whereas the 2D analysis included *n* = 625 control mitochondrial cross-sections and *n* = 212 patient-derived mitochondrial cross-sections. Group differences were assessed using a two-sided Mann–Whitney U test followed by Benjamini–Hochberg false discovery rate correction. Exact nominal *p* values and corrected *q* values are shown in Fig. 7c,d, and statistical outputs, including descriptive statistics, Mann–Whitney *U* statistics, and effect-size estimates, are provided with the analysis scripts.

**Figure 7.**
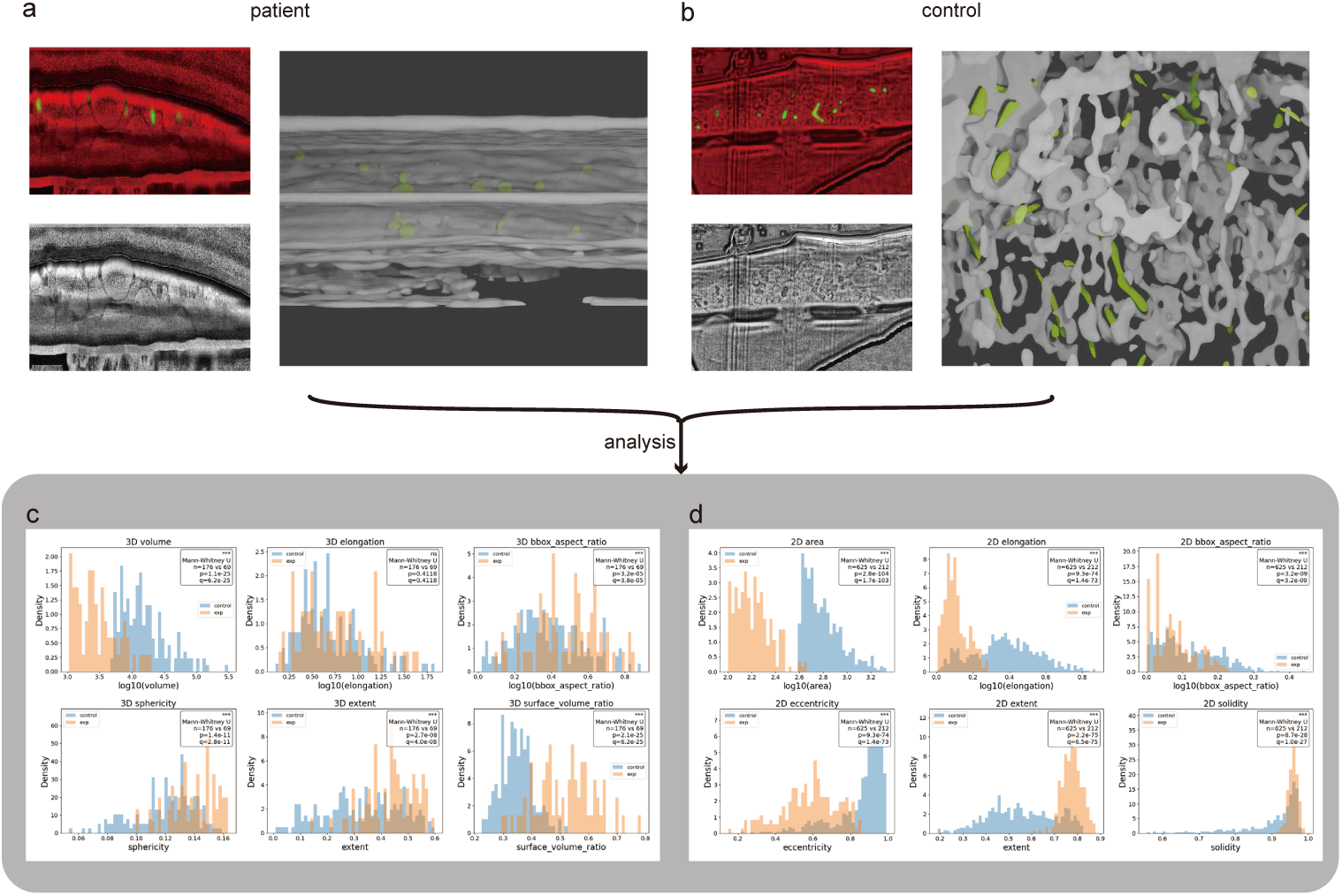
Quantitative morphometric analysis of mitochondria in control and Leigh syndrome patient cells. **a,b,** Representative two-dimensional slices showing raw images, SparseSeg predictions, and corresponding three-dimensional reconstructions. In the prediction overlays, red denotes background, green denotes predicted mitochondria, and blue denotes sparse annotation when available. In the three-dimensional renderings, white shows raw image context and green shows predicted mitochondria. **c,** Three-dimensional morphology distributions computed from reconstructed mitochondrial objects, including volume, elongation, bounding-box aspect ratio, sphericity, extent, and surface-to-volume ratio. The three-dimensional analysis included *n* = 176 control mitochondrial objects and *n* = 69 patient-derived mitochondrial objects. **d,** Two-dimensional morphology distributions computed from mitochondrial cross-sections, including area, elongation, bounding-box aspect ratio, eccentricity, extent, and solidity. The two-dimensional analysis included *n* = 625 control mitochondrial cross-sections and *n* = 212 patient-derived mitochondrial cross-sections. Histograms show density-normalized distributions. Each mitochondrial object or cross-section was measured once. Statistical significance was assessed using a two-sided Mann–Whitney U test, followed by Benjamini–Hochberg false discovery rate correction across morphology descriptors. Exact nominal *p* values and corrected *q* values are shown in each panel. These object-level measurements show reduced mitochondrial size and increased compactness and rounding in Leigh syndrome patient cells, consistent with mitochondrial fragmentation.

We next quantified three-dimensional mitochondrial morphology.

Three-dimensional morphometric measurements were computed directly from reconstructed mitochondrial volumes, including volume, elongation, bounding-box aspect ratio, sphericity, spatial extent, and surface-to-volume ratio. As shown in Fig. 7c, segmented mitochondria from the patient-derived example showed a shift toward smaller volumes, accompanied by increased extent and sphericity, consistent with a transition from elongated, interconnected networks to more compact and fragmented morphologies.

Notably, changes in elongation and bounding-box aspect ratio are less pronounced despite clear morphological differences. This can be attributed to the fact that mitochondria in control cells, although intrinsically elongated, often adopt curved and twisted configurations, such that the spatial regions they occupy are relatively compact and approximately isotropic at the bounding-box level. Consequently, even when mitochondria in patient cells undergo fragmentation or become more spherical, the overall spatial extent of individual structures remains comparable. As a result, elongation and bounding-box aspect ratio show limited variation, whereas the transition from elongated to more rounded morphologies is primarily reflected by increases in sphericity and extent.

We further observe an increase in the surface-to-volume ratio. In our implementation, this metric is estimated from surface voxels and therefore effectively measures the volume of a shell with finite thickness relative to the total volume. As mitochondrial volume decreases, the relative contribution of this shell increases, leading to an apparent elevation in the surface-to-volume ratio. Thus, this metric is sensitive to object size and may not be directly comparable across structures with substantially different volumes.

We also quantified two-dimensional mitochondrial morphology.

To assess whether disease-associated differences are also reflected at the slice level, we computed two-dimensional shape descriptors from individual cross sections, including area, elongation, eccentricity, extent, bounding-box aspect ratio, and solidity (Fig. 7d).

Consistent with the three-dimensional analysis, and in some cases more pronounced, these two-dimensional measurements indicate that slice-level analysis captures relevant morphological differences. Segmented mitochondria in patient cells exhibit increased solidity and extent, together with reduced eccentricity and elongation, consistent with more compact and rounded cross-sectional profiles. In addition, the decrease in area is indicative of mitochondrial fragmentation.

In contrast, changes in the bounding-box aspect ratio are less pronounced, consistent with the three-dimensional observations, suggesting that this metric is less sensitive for capturing morphological differences in this setting.

Finally, we examined biological consistency and implications.

The trends identified in our analysis are consistent with prior qualitative observations reported in the original study [12], and are here recovered automatically and quantitatively across the entire volume using model-derived segmentations (Fig. 7c,d). In particular, the observed differences in mitochondrial morphology, including fragmentation and increased rounding in Leigh syndrome patient cells, are in strong agreement with previous manual analyses.

These results demonstrate that previously reported cytoarchitectural alterations can be robustly reproduced using automated segmentation followed by large-scale morphometric analysis, without the need for manual tracing.

Together, these findings indicate that the output of our framework extends beyond visualization or pixel-wise segmentation accuracy and is sufficiently reliable to support statistically meaningful, high-throughput biological analysis in cryo-native FIB–SEM datasets. This capability enables unbiased, volume-wide quantification of organelle morphology and provides a scalable foundation for studying native-state cellular architecture in both healthy and disease-relevant contexts.

## Discussion

Recent deep learning approaches have achieved impressive results for conventional vEM segmentation [6, 14–16, 30], but many existing workflows still rely on densely annotated, high-contrast datasets and may require additional adaptation when applied to new imaging conditions or biological contexts.

A central contribution of this work is to formulate sparse organelle segmentation as a *target-conditioned discovery process*. Rather than treating segmentation only as the exhaustive identification of predefined organelle classes from densely annotated datasets, SparseSeg uses sparse, context-specific supervision to adapt segmentation to a given volume. This formulation is particularly useful for both cryo-vEM and conventional vEM imaging, where organelle appearance is strongly modulated by cellular environment, specimen preparation, and imaging conditions, and where dense voxel-level annotation is often impractical.

Our results show that models trained to recognize a specific organelle under one condition do not always generalize reliably to new datasets, even when overall organelle morphology is preserved. This observation does not imply that generalist segmentation models, domain adaptation methods, or foundation models cannot improve cross-domain generalization. Instead, SparseSeg addresses a complementary scenario: when users encounter a new vEM dataset, imaging condition, cell type, or target structure for which dense annotations are unavailable. In such cases, sparse dataset-specific exemplars provide a rapid and annotation-efficient way to adapt the model to local ultrastructural context and propagate target identity within the same volume.

Thus, target-conditioned discovery should be viewed as a practical complement to broadly trained segmentation models rather than a replacement for them. The refined labels generated by SparseSeg may also serve as pseudo-labels, initialization masks, or fine-tuning data for more general segmentation frameworks.

Recent progress in generalist vision foundation models and promptable segmentation models, including SAM-, MedSAM-, and SAM2-style architectures, provides an important opportunity for future extensions of SparseSeg [31–33]. Rather than competing with such models, SparseSeg could be integrated with them in several ways: sparse annotations could provide prompts or initialization masks, SparseSeg-refined predictions could serve as pseudo-labels for fine-tuning, and pretrained foundation features could improve robustness to heterogeneous image contrast. This integration may combine the broad visual priors of foundation models with the dataset-specific adaptability of sparse target-conditioned discovery.

By combining context-aware representation learning with geometry-consistent refinement, our framework bridges data-driven learning and explicit structural constraints. This hybrid strategy enables progressive expansion of reliable labels while suppressing spurious predictions, providing a practical mechanism for iterative self-improvement without sacrificing morphological fidelity. More broadly, these results suggest that scalable vEM analysis may benefit from complementing organelle-centric classifiers with adaptive, supervision-efficient systems that leverage limited expert input to discover structure within each dataset.

Beyond segmentation accuracy, the proposed framework enables automated, high-throughput quantitative analysis of native-state cellular architecture. In disease-relevant cryo-FIB–SEM datasets, such as Leigh syndrome patient cells, three-dimensional morphometric analysis reveals systematic shifts in mitochondrial volume, elongation, and sphericity, consistent with fragmentation and loss of network integrity. While such trends have been qualitatively reported in prior studies [12], our approach enables their recovery in an automated and scalable manner. This capability highlights the importance of accurate volumetric reconstruction for linking ultrastructure to cellular function and pathology, and is consistent with observations in other native-state systems, including mineralized tissues and muscle ultrastructure, where subtle three-dimensional organization encodes critical biological function [34–36].

Several limitations and potential failure modes should be noted. First, SparseSeg depends on the quality and representativeness of the initial sparse annotations. If the user-provided ROIs cover only a narrow subset of the target morphology, the model may preferentially propagate similar-looking regions while missing target instances with different orientations, sizes, or local contrast. Conversely, if the initial annotations accidentally include ambiguous boundaries or nearby non-target structures, these errors may be reinforced during iterative label expansion. Second, although the geometry-consistent refinement step is designed to suppress spurious predictions, it can also introduce failure modes. Overly permissive refinement thresholds may allow false-positive regions with similar shape statistics to be added as pseudo-labels, leading to error propagation in subsequent iterations. In contrast, overly strict shape or boundary–region filtering may remove true target structures with atypical morphology, causing false negatives. This trade-off is illustrated in the failure-case analysis in Supplementary Fig. 4, where different sparse annotation choices lead to different prediction coverage and error patterns. Third, SparseSeg may struggle when the target structure has weak contrast, highly fragmented boundaries, or strong morphological similarity to neighboring ultrastructures. In such cases, the model may confuse the target with visually similar organelles or background membranes, particularly when only very few positive ROIs are provided. Last, the current implementation is target-specific and segments one user-defined structure at a time. Therefore, simultaneous segmentation of multiple ultrastructures, such as mitochondria, vacuoles, filaments, ER, or Golgi apparatus, currently requires separate sparse annotations and separate target-specific training or inference runs. This is an important limitation of the present framework. At the same time, this design reflects the goal of SparseSeg as a target-conditioned discovery method rather than a closed-set multi-class segmentation model. The target-specific formulation allows rapid adaptation to new structures and imaging contexts from minimal user-provided exemplars, but it does not yet provide a unified multi-target segmentation model.

These limitations indicate that SparseSeg should be used as an interactive, target-conditioned segmentation tool rather than a fully automatic universal or simultaneous multi-class organelle classifier. In practice, performance can be improved by providing more representative sparse annotations, inspecting intermediate predictions, and adjusting refinement thresholds to balance false-positive suppression and recall. Future extensions could address the single-target limitation by using multi-channel sparse annotations, a shared backbone with target-specific output heads, or target embeddings that allow multiple ultrastructural classes to be segmented within the same volume.

Additional future directions include integration with correlative light and electron microscopy to directly link functional imaging with native ultrastructure [37, 38], extension to large tissue volumes and community-driven annotation platforms [39, 40], incorporation of self-supervised or generative learning strategies [41–43], and development of multi-target extensions that can jointly model multiple user-defined ultrastructures within the same volume. Together, these developments position cryo-vEM as a quantitative and hypothesis-generating platform for studying cellular organization in health and disease.

## Methods

Our framework implements a target-conditioned, sparsity-driven segmentation paradigm through three tightly integrated modules: (i) target-conditioned sparse supervision, (ii) context- and geometry-aware representation learning, and (iii) iterative discovery and refinement. Together, these components enable robust cryo-volume electron microscopy (cryo-volume EM) segmentation under extreme annotation sparsity and domain shift. We denote the aligned input image volume as *V*, the sparse positive label mask as *M*, and the optional negative annotation mask as *N*. Detailed implementation parameters, preprocessing scripts, training scripts, inference scripts, refinement scripts, and example configuration files are provided in the Supplementary Methods and in the repository README.

### Module I: Target-conditioned sparse supervision

All volumetric electron microscopy (vEM) datasets used in this study were prepro-cessed and annotated using a custom Fiji/ImageJ macro to standardize geometry, correct inter-slice misalignment, and generate sparse supervision signals. This step is not specific to our implementation and can be replaced by other appropriate preprocessing pipelines.

The volume is aligned using a SIFT-based translation correction algorithm implemented in ImageJ to compensate for inter-slice drift and minor stage instability. This step can be omitted if the dataset is already well aligned. The resulting aligned volume corresponds to the input image volume *V*.

Rather than requiring dense voxel-level annotations, SparseSeg relies on the sparse positive label mask *M* to condition the segmentation task. In practice, expert users annotate a small number of representative regions of interest (ROIs) corresponding to the target organelle, and these ROIs are converted into the binary sparse positive label mask *M*.

When available, user-provided negative ROIs indicating regions where the target structure is confidently absent are converted into the optional binary negative annotation mask *N*. Together, the pair (*M, N*) defines the target-conditioned supervision signal that enables the model to adapt to dataset-specific context, imaging conditions, and annotation strategies.

All outputs, including the aligned image volume *V*, the sparse positive label mask *M*, and the optional negative annotation mask *N*, are stored as TIFF stacks and serve as the inputs for downstream learning.

### Module II: Context- and geometry-aware representation learning

#### Sparse sampling and training dataset construction

We construct a feature-enhanced representation of the input volume together with a set of complementary supervision signals, including boundary maps, negative masks, and feature-based reference maps. Based on these components, training samples are generated using a patch-based sampling strategy centered on sparsely annotated regions, with a small proportion of background sampling to improve robustness. Each sample is formulated as a multi-channel patch that integrates both image features and supervision signals, providing a unified input for subsequent model training (see Supplementary).

This sampling strategy enables efficient construction of informative training data from extremely sparse annotations, while preserving structural context and boundary information that are critical for accurate segmentation.

#### Multi-scale representation learning and network architecture

Segmentation is performed using a customized Multi-Kernel U-Net architecture. Each convolutional block consists of three parallel branches with kernel sizes 3, 5, and 7, followed by batch normalization and ReLU activation. Branch outputs are concatenated and fused through a 1 × 1 convolution, enabling simultaneous encoding of fine membrane boundaries, intermediate organelle morphology, and broader contextual structures.

The encoder comprises three resolution levels with feature dimensions 64, 128, and 256, interleaved with 2 × 2 max pooling. The decoder mirrors this structure using transposed convolutions and skip connections. Although attention-gated skip connections were implemented, ungated connections were used in the reported experiments for improved training stability. A final 1 × 1 convolution produces slice-wise probability maps, which are assembled across *z*-slices to form the final three-dimensional prediction volume.

#### Geometry-aware learning objective

To accommodate sparse and incomplete annotations while enforcing structural coherence, we employ a composite geometry-aware loss. The network predicts two probability maps, corresponding to the region-interior channel and the boundary/edge channel. In the current implementation, the masked binary cross-entropy (BCE) loss provides the primary sparse-supervision signal, while the remaining terms are used as weak auxiliary regularizers to improve structural coherence and suppress spurious predictions.

The total loss is written in a coarse form as

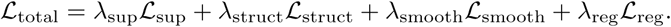

Here, ℒ_sup_ denotes the supervised masked BCE loss applied to the region-interior and boundary/edge prediction channels. The region-interior channel is supervised by the sparse positive annotation mask, whereas the boundary/edge channel is supervised by the boundary mask derived from the same sparse annotation. ℒ_struct_ denotes auxiliary structural losses, including region-consistency and region-contrast terms computed from feature-based reference maps. ℒ_smooth_ denotes an image-guided smoothness loss that reduces fragmented activations while preserving image-defined boundaries. ℒ_reg_ denotes regularization terms that suppress spurious activations and promote compact predictions. Detailed definitions of each loss component are provided in Supplementary.

In the reported experiments, the supervised masked BCE term was assigned the dominant weight, whereas the structural, smoothness, and regularization terms were assigned substantially smaller weights. Specifically, the default weights were *λ*_sup_ = 10, *λ*_struct_ = 0.1, *λ*_smooth_ = 0.1, and *λ*_reg_ = 0.05. Our ablation analysis shows that the masked BCE loss is essential for learning meaningful target segmentation, whereas the auxiliary losses have weaker effects in the mitochondria benchmark. Therefore, these additional terms should be interpreted as optional regularizers rather than equally important primary objectives. They are retained to support more challenging settings, such as low-contrast images, noisy sparse annotations, or targets with weak boundaries, but can be reduced or disabled when they do not improve validation performance. In practice, training stability is maintained by assigning small weights to the auxiliary losses and monitoring individual loss components during optimization. Training is performed using the Adam optimizer with a learning rate of 1 × 10*^−^*^3^, batch size of 12, and 60 epochs per iteration.

### Module III: Iterative discovery and refinement

#### Geometry-consistent label refinement

Raw network predictions may contain fragmented or spurious regions, particularly under low contrast. To enable reliable target discovery and prevent error accumulation, we apply a geometry-consistent refinement stage.

Predicted masks are decomposed into region interiors and boundary-like components. Connected components are extracted and evaluated using shape descriptors, including circularity, aspect ratio, extent, solidity, eccentricity, Euler number, and low-order Hu invariant moments. Only components whose shape statistics are consistent with sparse reference annotations and satisfy biologically plausible size and topology constraints are retained.

#### Boundary–region association and label propagation

To enforce contour–area consistency, refined region candidates are further validated using predicted boundary structures. Binary edge and area masks are matched via bounding-box intersection over union (IoU) calculations. Region components are preserved only if they exhibit sufficient spatial overlap with thin boundary structures, suppressing blob-like artifacts.

Validated predictions are merged with existing annotations to form an expanded supervision set. These augmented labels are fed back into the training pipeline in subsequent iterations. Through this iterative discovery-and-refinement process, the model progressively expands supervision coverage, improves segmentation quality, and remains robust to domain shift and annotation sparsity.

## Data Availability

All volumetric electron microscopy datasets analyzed in this study are publicly available. The Janelia COSEM datasets can be accessed through the OpenOrganelle data portal and the Janelia COSEM data repository. The EMPIAR datasets used in this study are available from the Electron Microscopy Public Image Archive under accession numbers EMPIAR-10311, EMPIAR-10442, EMPIAR-10392, EMPIAR-10515, EMPIAR-11415, EMPIAR-11416, EMPIAR-11417, EMPIAR-11419, and EMPIAR-11420.

Example data for running SparseSeg, including representative input volumes, sparse annotations, and example segmentation outputs, are available in the SparseSeg GitHub repository: https://github.com/s616442197-dotcom/SparseSeg/tree/master.

The morphometric and benchmarking results reported in this study were generated from the publicly available datasets listed above using the analysis scripts provided in the code repository. No restrictions apply to the publicly available datasets beyond the access and reuse terms specified by the original data repositories.

## Code Availability

The complete SparseSeg codebase, including the segmentation framework, training scripts, inference scripts, benchmarking scripts, morphometric analysis scripts, example sparse annotations, default parameter settings, and documentation, is available as open-source software at: https://github.com/s616442197-dotcom/SparseSeg/tree/master.

## Acknowledgements

This work was supported by the National Natural Science Foundation of China (grant no. 12504242); the Fundamental and Interdisciplinary Disciplines Breakthrough Plan of the Ministry of Education of China (grant no. JYB2025XDXM502); the National Key R&D Program of China (grant no. 2025YFA0923100); the Fundamental Research Funds for the Central Universities (grant no. 226-2025-00252).

## Author Contributions

Y.Z. conceived the project. Y.Z. and Q.O. supervised this study. B.S. developed the SparseSeg model, implemented the segmentation pipeline, performed computational experiments and analyses. Y.L. curated, cleaned and organized the volume electron microscopy datasets used in this study. B.S and Y.Z. wrote the manuscript with input from all the authors.

## Competing Interests

The authors declare that they have no competing interests.

## Notes

### Competing Interest Statement

The authors have declared no competing interest.

