## Supplementary for "SparseSeg: Target-Conditioned Discovery Segmentation of Cryo-Volume Electron Microscopy Under Sparse Annotation"

Supplementary Information

Bowen Shi, Yanjun Li, Qi Ouyang, and Yanan Zhu

#### Supplementary Figures

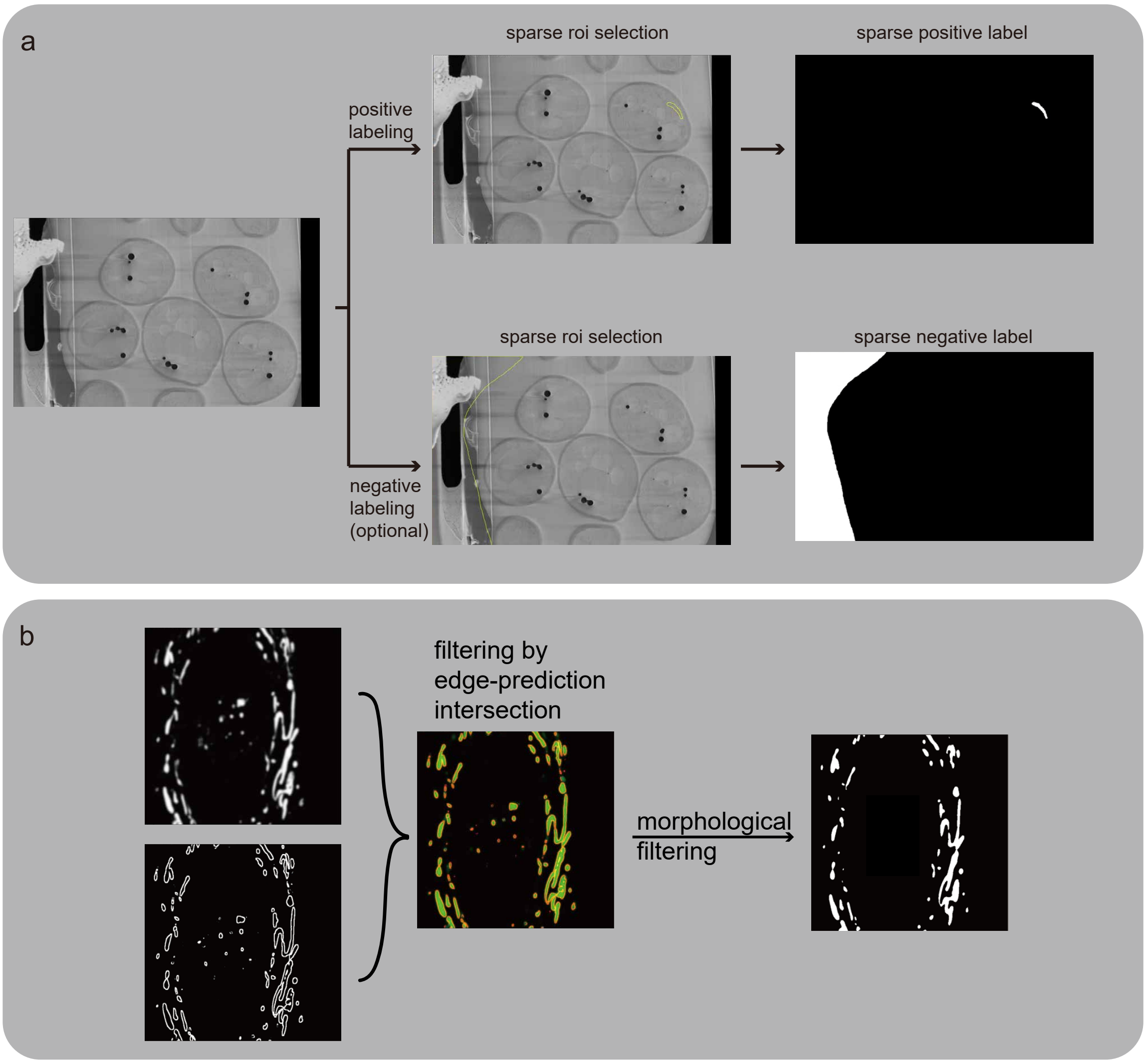


**Supplementary Fig. 1.** Preprocessing and refinement of raw volume electron microscopy (vEM) data. **a,** Sparse positive target annotations and explicit background labeling used to initialize supervised training under weak annotation. **b,** Morphological and topology-aware post-processing applied to model predictions, including shape-based filtering and connectivity refinement, to suppress spurious detections and recover coherent three-dimensional structures.

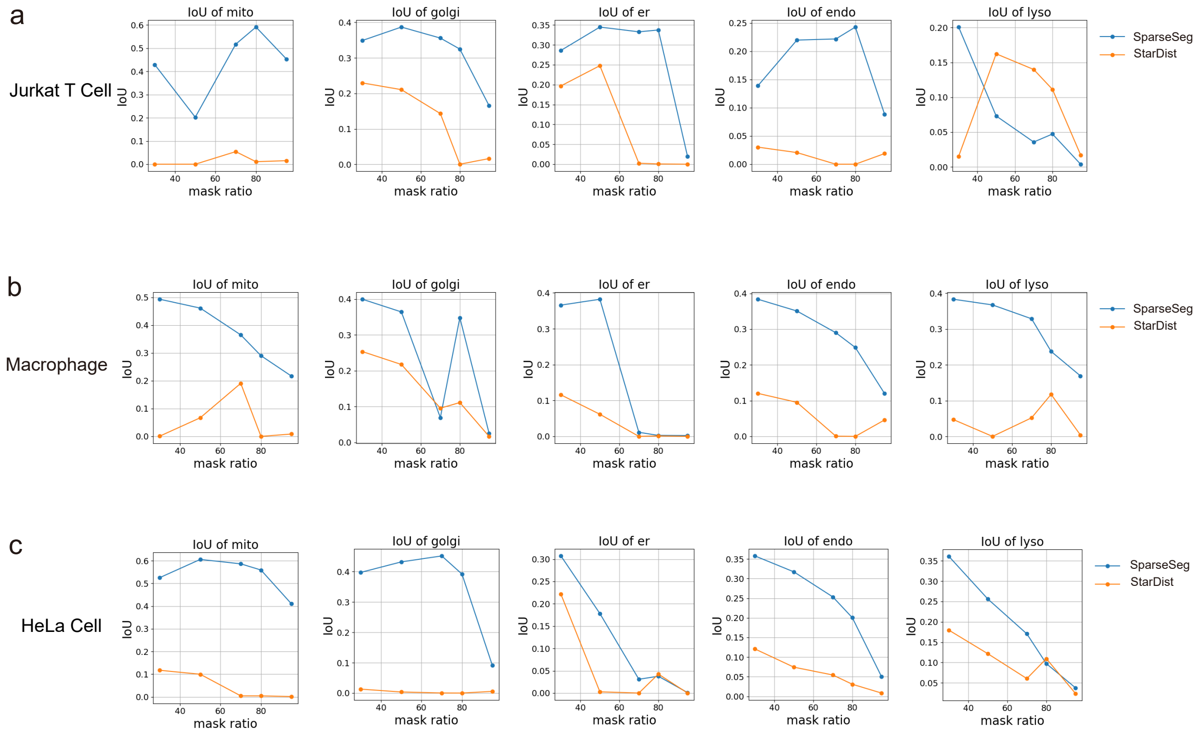


**Supplementary Fig. 2.** Additional benchmarking results comparing SparseSeg and StarDist across multiple organelles and cell types under varying levels of label sparsity. **a,** Jurkat T cells. **b,** Macrophages. **c,** HeLa cells. Performance differences across organelles reflect both model behavior and variability in annotation quality.

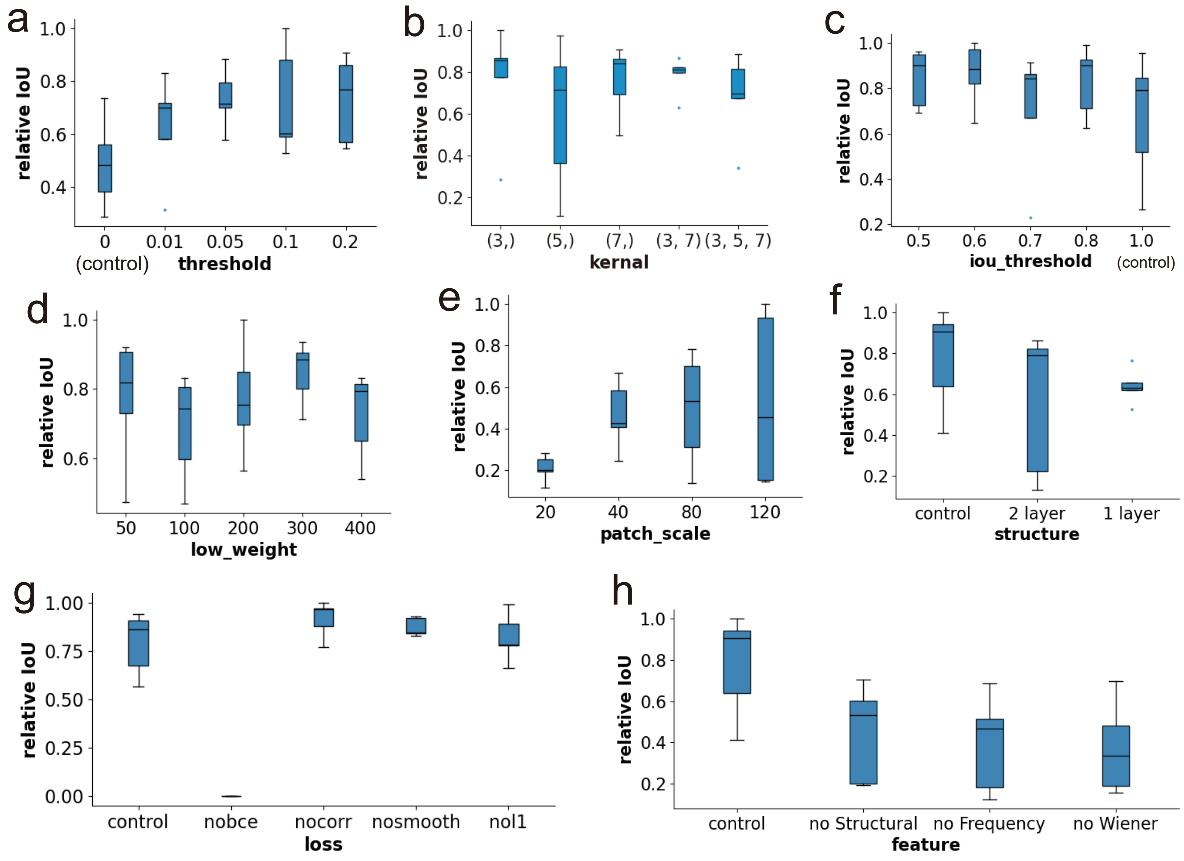


**Supplementary Fig. 3.** Ablation and robustness analysis of SparseSeg. **a,** Effect of the threshold used for shape refinement. **b,** Effect of kernel size and the number of kernels. **c,** Effect of the IoU threshold used for boundary–region filtering. **d,** Effect of the low-weight coefficient used to down-weight regions far from positive annotations. **e,** Effect of patch size on segmentation performance. **f,** Ablation of the U-Net backbone structure. **g,** Ablation of different loss components. **h,** Ablation of feature groups, including structural filters, frequency filters, and Wiener-denoised features.

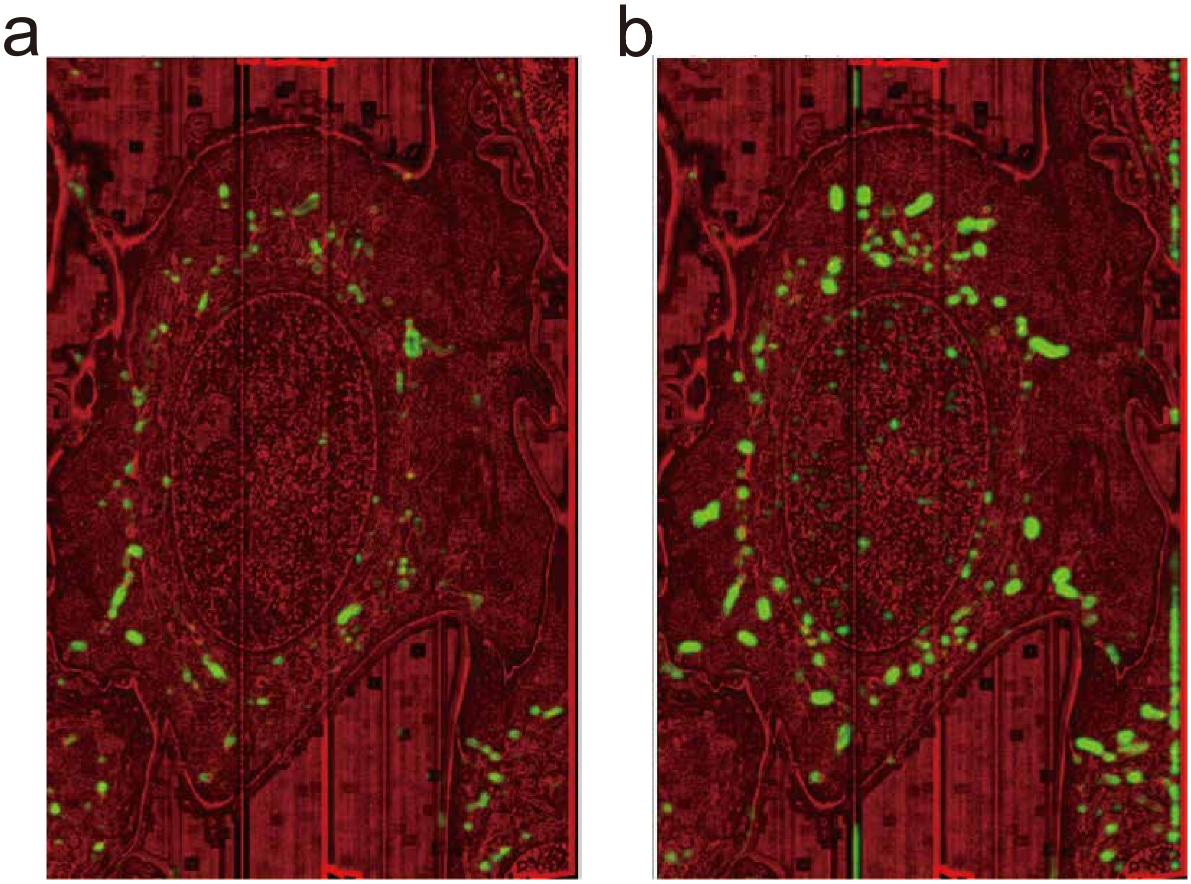


**Supplementary Fig. 4.** Prediction variability caused by different sparse positive annotations. **a,** Prediction from trial 1. **b,** Prediction from trial 3. Compared with trial 1, trial 3 identifies more candidate target structures but also introduces more false-positive predictions.

### 8 Supplementary Tables

**Supplementary Table 1.** CPU and memory usage records from five representative SparseSeg trials. For process-level CPU utilization, 100% corresponds to one CPU core; therefore, values above 100% indicate multi-core CPU usage.

| Trial | CPU avg (%) | CPU max (%) | Proc mem avg (GB) | Proc mem max (GB) | Sys mem avg (%) | Sys mem max (%) |
|---|---|---|---|---|---|---|
| 1 | 752.80 | 3035.80 | 14.484 | 29.753 | 32.18 | 56.60 |
| 2 | 844.45 | 3049.80 | 14.242 | 29.795 | 31.85 | 56.60 |
| 3 | 867.04 | 3048.60 | 14.267 | 29.770 | 31.81 | 56.40 |
| 4 | 863.74 | 3046.30 | 14.255 | 29.967 | 31.78 | 56.80 |
| 5 | 795.04 | 3078.40 | 14.330 | 29.889 | 31.86 | 56.50 |

**Supplementary Table 2.** GPU utilization and GPU memory usage records from five representative SparseSeg trials. GPU memory usage was stable across runs, with approximately 4.1 GB used out of 16.3 GB available GPU memory.

| Trial | GPU util avg (%) | GPU util max (%) | GPU mem avg (MB) | GPU mem max (MB) | GPU total (MB) |
|---|---|---|---|---|---|
| 1 | 19.74 | 64 | 4109.00 | 4109.00 | 16303 |
| 2 | 22.09 | 64 | 4109.00 | 4109.00 | 16303 |
| 3 | 22.74 | 65 | 4112.97 | 4120.00 | 16303 |
| 4 | 23.13 | 64 | 4118.79 | 4125.00 | 16303 |
| 5 | 20.67 | 95 | 4105.00 | 4105.00 | 16303 |

### Supplementary Notes

#### Ethics approval and consent to participate

This study exclusively used publicly available imaging datasets. No human subjects or animals were directly involved in this research. All original data acquisition protocols are described in the corresponding primary publications.

#### Consent for publication

Not applicable.

#### Materials availability

No new biological materials were generated in this study.

### Supplementary Methods

#### Datasets

The training and evaluation datasets used in this study comprise multiple volumetric electron microscopy datasets obtained from two public resources: the Janelia COSEM whole-cell FIB–SEM atlas and the Electron Microscopy Public Image Archive (EMPIAR).

##### Janelia COSEM datasets (mammalian single cells)

Three densely annotated mammalian single-cell datasets were obtained from the Janelia COSEM project. All datasets were imaged using focused ion beam scanning electron microscopy (FIB–SEM) following high-pressure freezing and freeze-substitution resin embedding, and are accompanied by expert manual organelle segmentations provided by the COSEM Project Team.

The first dataset corresponds to a wild-type interphase HeLa cell (ATCC CCL-2). Cells were prepared by high-pressure freezing, followed by freeze substitution in acetone containing 2% osmium tetroxide ($OsO_4$), 0.1% uranyl acetate, and 3% $H_2O$, and subsequently embedded in Eponate 12 resin. FIB–SEM imaging and post-processing were performed at HHMI Janelia. Dense manual segmentations of major membrane-bound organelles were generated by the COSEM Project Team and used as ground-truth annotations in this study.

The second dataset consists of a wild-type THP-1–derived macrophage (ATCC TIB-202). Cells were differentiated using phorbol 12-myristate 13-acetate (PMA) and prepared using the same high-pressure freezing and freeze-substitution protocol as the HeLa dataset. Imaging, post-processing, and comprehensive organelle segmentation were performed by the Janelia COSEM team.

The third dataset comprises wild-type Jurkat T cells (clone E6-1, ATCC TIB-152). Cells were prepared and imaged using identical cryo-FIB/SEM workflows, including high-pressure freezing and freeze substitution. Expert manual segmentations provided by the COSEM Project Team were used as reference annotations.

Together, these three datasets provide whole-cell, high-resolution volumetric EM data with dense ground-truth annotations for major membrane-bound organelles, and serve as the primary source of supervised labels for training and benchmarking in this study.

##### EMPIAR datasets (diverse organisms and tissues)

To increase biological diversity and assess model generalization across organisms, tissues, and imaging modalities, six additional volumetric EM datasets were obtained from the Electron Microscopy Public Image Archive (EMPIAR).

EMPIAR-10311 consists of a FIB–SEM volume of an interphase HeLa cell, providing an additional mammalian single-cell dataset distinct from the COSEM atlas [1].

EMPIAR-10442 contains serial block-face scanning electron microscopy (SBF–SEM) data of the *Arabidopsis thaliana* root phloem unloading zone, representing plant tissue with highly specialized cellular organization [2].

EMPIAR-10392 comprises FIB–SEM data capturing *Plasmodium falciparum* across cytokinesis, introducing a unicellular eukaryotic parasite with markedly different ultrastructural features [3].

EMPIAR-10515 provides serial cryo-FIB/SEM datasets of Leigh syndrome patient cells and matched control cells, enabling disease-relevant quantitative analysis under cryo-native conditions [4].

EMPIAR-11415 Cryo serial FIB SEM of mouse brain tissue [5].

EMPIAR-11416 consists of cryo serial FIB–SEM data of *Saccharomyces cerevisiae*, representing a unicellular eukaryotic model organism with compact cellular architecture [5].

EMPIAR-11417 Cryo serial FIB/SEM of Vero cells [5].

EMPIAR-11419 Cryo serial FIB/SEM of HeLa cells [5].

EMPIAR-11420 comprises cryo serial FIB–SEM data of mouse heart tissue, representing multicellular tissue-scale organization with densely packed and highly ordered ultrastructure [5].

Collectively, the EMPIAR datasets span mammalian cells, plant tissue, unicellular eukaryotes, and multicellular tissue, introducing substantial variation in ultrastructural organization, organelle morphology, and image contrast. These datasets enable rigorous evaluation of model robustness and generalization across biological systems and imaging conditions.

#### Detailed dataset construction and feature extraction

This Supplementary provides a detailed description of the dataset construction and feature extraction procedures used for training and validation. These design choices were motivated by the unique characteristics of cryo-native volume electron microscopy (EM) data, including low contrast, subtle membrane signals, and extremely sparse annotations.

**Multi-scale feature embedding from volumetric data** To enhance the representation of volumetric EM data, we construct a multi-channel feature volume from the input image volume $V \in \mathbb{R}^{D\times H\times W}$. This feature-extraction step is used as a conventional image preprocessing and input-enhancement procedure, rather than as a newly proposed feature-extraction algorithm. The resulting feature volume is denoted as $F \in \mathbb{R}^{D\times C\times H\times W}$, where $C$ is the number of extracted feature channels.

For each slice $I \in \mathbb{R}^{H\times W}$, we compute a set of standard image-processing descriptors, including local intensity statistics, multi-scale filtering responses, and frequency-domain representations. These descriptors are intended to improve the visibility of low-contrast vEM structures and provide structure-aware inputs for sparse target-conditioned learning.

Local statistics are computed using a sliding window (size $5 \times 5$), including the local mean $\mu$, variance $\sigma^2$, and gradient magnitude:

$$\mu = \text{UniformFilter}(I), \quad \sigma^2 = \text{UniformFilter}(I^2) - \mu^2, \tag{1}$$

$$|\nabla I| = \sqrt{\left(\frac{\partial I}{\partial x}\right)^2 + \left(\frac{\partial I}{\partial y}\right)^2}. \tag{2}$$

Multi-scale structural features are extracted by applying Gaussian smoothing at multiple scales ($\sigma \in \{3, 5, 7\}$), followed by computation of Hessian eigenvalues and a Frangi-like vesselness measure to capture curvilinear structures. In addition, Laplacian-of-Gaussian (LoG) responses are included to emphasize blob-like structures.

Frequency-domain features are computed via Fourier filtering using radial masks, including low-pass, high-pass, and band-pass components, capturing complementary global structural information. Wiener filtering with multiple kernel sizes is further applied to suppress noise while preserving structural details.

These operations are used jointly as complementary input channels. We do not attribute the final segmentation performance to any single handcrafted feature channel; instead, the main methodological contribution of SparseSeg lies in the target-conditioned sparse-supervision framework, the segmentation backbone, and the geometry-consistent iterative refinement pipeline.

All feature maps are stacked along the channel dimension to form a feature tensor:

$$\phi(I) \in \mathbb{R}^{C_0 \times H \times W}, \tag{3}$$

where $C_0$ denotes the number of two-dimensional feature channels.

To incorporate three-dimensional context, for each slice index $z$, we construct a local stack:

$$X_z = V[z - t : z + t + 1], \tag{4}$$

where $t$ denotes the axial half-thickness.

Two-dimensional features are extracted independently for each slice within the stack:

$$\{\phi(X_z^{(k)})\}_{k=1}^{2t+1}. \tag{5}$$

These features are aggregated along the axial dimension using statistical pooling:

$$\phi_{\text{mean}} = \text{mean}_k\, \phi(X_z^{(k)}), \quad \phi_{\text{var}} = \text{var}_k\, \phi(X_z^{(k)}). \tag{6}$$

The aggregated feature representation is obtained by concatenation:

$$f_z = \text{concat}(\phi_{\text{mean}}, \phi_{\text{var}}), \quad f_z \in \mathbb{R}^{C \times H \times W}, \tag{7}$$

where $C = 2C_0$.

The aggregated feature map is assigned to the corresponding slice:

$$F[z] = f_z. \tag{8}$$

Repeating this process for all valid slice indices yields the final feature volume $F$, which preserves spatial resolution while incorporating local three-dimensional contextual information.

**Boundary mask construction** To explicitly supervise thin membrane structures and boundary regions, we derive an edge map from the foreground mask $M$ using morphological operations.

Let $M \in \{0,1\}^{H \times W}$ denote the binary foreground mask. We first compute its dilated and eroded versions using morphological filtering:

$$M_{\text{dil}} = \text{Dilate}(M), \quad M_{\text{ero}} = \text{Erode}(M), \tag{9}$$

where dilation and erosion are implemented via max-pooling and its dual operation with a predefined kernel size.

The boundary map is then defined as the symmetric difference between the dilated and eroded masks:

$$E = M_{\text{dil}} \oplus M_{\text{ero}}, \tag{10}$$

where $\oplus$ denotes the logical exclusive-or (XOR) operation.

This formulation produces a thin, one-pixel-wide boundary approximation that captures transitions between foreground and background regions. Compared to gradient-based edge detection, this approach is directly derived from supervision masks and is therefore robust to noise and intensity variations in cryo-vEM data.

The resulting edge map $E$ is used as an additional supervision signal to explicitly guide the model in learning boundary localization and preserving fine structural details.

**Hard and soft negative mask construction** To refine supervision signals, we construct both hard and soft negative masks based on the foreground mask $M$. Morphological operations are first applied to $M$ to generate boundary regions and surrounding negative rings $N_M$, analogous to the construction of the edge map $E$. These are combined with user-provided negative annotations $N$ to form a hard negative mask:

$$N_f = N_M \cup N, \tag{11}$$

which integrates both explicit supervision and geometry-derived structural constraints.

To further suppress spurious positive predictions in distant background regions, we construct a soft-negative mask $S$ based on the distance transform of $M$. Specifically, we compute:

$$D = \mathrm{dist}(1 - M), \tag{12}$$

where $D$ denotes the Euclidean distance to the nearest positive voxel.

A distance-based sigmoid mapping is then applied:

$$S = \alpha \cdot \max\left(0, \frac{1}{1 + \exp\left(-\frac{D-R}{k}\right)} - \epsilon\right), \tag{13}$$

where $R$ controls the transition radius, $k = R/6$ determines the smoothness of the transition, $\epsilon$ suppresses near-boundary responses, and $\alpha$ is a scaling factor.

This formulation assigns low weights to regions proximal to annotated positives and progressively increases penalties for voxels located farther away. Consequently, $S$ encourages the model to treat distant background regions as negative while maintaining a smooth transition near object boundaries.

**Reference map construction via feature projection** To enhance the discriminative representation of target structures, we construct reference maps by projecting feature volumes onto supervision-guided directions.

Given a feature volume $F \in \mathbb{R}^{D\times C\times H\times W}$, we compute mean feature vectors over positive and negative regions. Specifically, positive regions are defined either by boundary annotations $E$ or area annotations $M$, while negative regions are defined by the hard negative mask $N_f$. The corresponding mean feature vectors are given by:

$$\boldsymbol{\mu}_{\text{pos}} = \frac{1}{|\Omega_{\text{pos}}|} \sum_{i\in\Omega_{\text{pos}}} \mathbf{x}_i, \quad \boldsymbol{\mu}_{\text{neg}} = \frac{1}{|\Omega_{\text{neg}}|} \sum_{i\in\Omega_{\text{neg}}} \mathbf{x}_i. \tag{14}$$

A projection direction is then defined as:

$$\mathbf{w} = \frac{\boldsymbol{\mu}_{\text{pos}} - \boldsymbol{\mu}_{\text{neg}}}{\|\boldsymbol{\mu}_{\text{pos}} - \boldsymbol{\mu}_{\text{neg}}\|_2 + \epsilon}. \tag{15}$$

The reference map is obtained by projecting the feature volume onto $\mathbf{w}$:

$$R(d, h, w) = \sum_{c=1}^{C} F(d, c, h, w) \cdot w_c. \tag{16}$$

This formulation yields scalar volumetric maps corresponding to different supervision signals, including edge-based reference maps $R_E \in \mathbb{R}^{D\times H\times W}$ and area-based reference maps $R_A \in \mathbb{R}^{D\times H\times W}$. These reference maps are used in subsequent loss computation to guide learning toward target-specific feature directions.

**Construction of training dataset** Training data are constructed using a patch-based sampling strategy from the volumetric image $V \in \mathbb{R}^{D\times H\times W}$. Sparse supervision is provided in the form of $M$ and $N$.

Candidate patch centers are first selected from all voxels satisfying $M > 0$, ensuring that each sampled patch contains target-associated signal. To improve background discrimination and suppress false-positive predictions, an additional small fraction of sampling centers is drawn from voxels satisfying $(N > 0)$. The number of such background-anchored samples is kept substantially smaller than that of positive samples to maintain supervision balance while improving robustness.

All candidate sampling centers are filtered to ensure that extracted patches remain fully within valid image boundaries. For a spatial patch size $(h, w)$ and axial half-thickness $t$, a valid sampling center $(z_m, x_m, y_m)$ must satisfy

$$t \le z_m < D - t, \quad \frac{h}{2} \le x_m < H - \frac{h}{2}, \quad \frac{w}{2} \le y_m < W - \frac{w}{2}. \tag{17}$$

Here, $z_m$ denotes the selected central slice, while the axial context around $z_m$ has already been incorporated into the feature volume during feature construction.

For each valid sampling center $(z_m, x_m, y_m)$, a local spatial patch is extracted from the centered region

$$\Omega_m = \left[x_m - \frac{h}{2} : x_m + \frac{h}{2},\ y_m - \frac{w}{2} : y_m + \frac{w}{2}\right]. \tag{18}$$

The input feature tensor for this training sample is therefore defined as

$$F_m = F\left[z_m,\ :,\ x_m - \frac{h}{2} : x_m + \frac{h}{2},\ y_m - \frac{w}{2} : y_m + \frac{w}{2}\right], \tag{19}$$

where $F \in \mathbb{R}^{D\times C\times H\times W}$ is the feature volume.

The corresponding supervision components are cropped from the same spatial region on slice $z_m$:

$$M_m,\ E_m,\ N_{f,m},\ S_m,\ R_{E,m},\ R_{A,m}. \tag{20}$$

These components are obtained by applying the same centered crop $\Omega_m$ to the positive mask, edge mask, hard negative mask, soft negative mask, edge-based reference map, and area-based reference map, respectively.

Together, these cropped image features and supervision signals define one training sample. The full training dataset is implemented using standard PyTorch data pipelines to enable efficient batch sampling and model optimization.

#### Network architecture

The segmentation model is based on a U-Net-like encoder–decoder architecture with symmetric skip connections, augmented by multi-kernel convolutional blocks to enhance multi-scale feature representation. The overall design aims to capture fine membrane boundaries, intermediate organelle morphology, and long-range contextual cues within a unified framework.

**Multi-kernel convolutional block.** Each encoder and decoder stage is implemented using a *multi-kernel convolutional block*. Given an input feature map $x \in \mathbb{R}^{C\times H\times W}$, the block applies multiple convolutional branches in parallel, with kernel sizes

$$\mathcal{K} = \{3,\ 5,\ 7\}. \tag{21}$$

Each branch consists of a $k \times k$ convolution (with appropriate zero padding), followed by batch normalization and a ReLU activation. The outputs of all branches are concatenated along the channel dimension and fused using a $1 \times 1$ convolution, followed by batch normalization and ReLU:

$$\mathrm{MKConv}(x) = \phi(\mathrm{BN}(W_{1\times1}[\phi(\mathrm{BN}(W_3 * x)) \parallel \phi(\mathrm{BN}(W_5 * x)) \parallel \phi(\mathrm{BN}(W_7 * x))])) \,, \tag{22}$$

where $\phi(\cdot)$ denotes the ReLU activation and $\parallel$ denotes channel-wise concatenation.

This design enables simultaneous extraction of local edge features, mid-scale structural patterns, and broader contextual information, which is particularly important for low-contrast cryo-volume EM data.

**Encoder.** The encoder consists of three resolution levels. At each level, a multi-kernel convolutional block increases the feature dimensionality, followed by $2 \times 2$ max-pooling for spatial downsampling. Specifically, the encoder stages map

$$C \to 64 \to 128 \to 256 \tag{23}$$

channels while progressively reducing spatial resolution by a factor of two at each level.

**Decoder.** The decoder mirrors the encoder structure. Feature maps are upsampled using transposed convolutions with stride 2, followed by concatenation with the corresponding encoder features via skip connections. Each concatenated feature map is processed by a multi-kernel convolutional block to refine spatial details and recover fine structures.

Although attention gates are implemented for skip connection filtering, they are disabled in the current experiments, and skip connections directly propagate encoder features to the decoder.

**Output layer.** The final decoder output is mapped to voxel-wise prediction logits using a $1 \times 1$ convolution. Depending on the task configuration, the output may contain one or multiple channels corresponding to region and edge predictions. A sigmoid activation is applied during training and inference to obtain probabilistic segmentation masks.

**Design rationale.** Compared to standard U-Net architectures, the proposed model introduces multi-kernel feature extraction at every resolution level. This design is intended to improve flexibility across cryo-volume EM targets with different sizes, shapes, and contrast characteristics. By avoiding a single fixed receptive field, the network can integrate information across multiple spatial scales, but multi-kernel configurations are not expected to guarantee the best performance for every individual segmentation task.

#### Loss formulation

**Notation.** For each sampled two-dimensional training patch, let $I \in \mathbb{R}^{H\times W}$ denote the corresponding input image or reference map. The network outputs two logit maps:

$$\hat{Y}_A, \hat{Y}_E \in \mathbb{R}^{H\times W}, \tag{24}$$

where $\hat{Y}_A$ denotes the area/interior logit map and $\hat{Y}_E$ denotes the edge logit map. The corresponding probability maps are

$$P_A = \sigma(\hat{Y}_A), \qquad P_E = \sigma(\hat{Y}_E). \tag{25}$$

Sparse positive area labels are denoted by $Y_A \in \{0,1\}^{H\times W}$, and edge labels derived from the sparse annotation are denoted by $Y_E \in \{0,1\}^{H\times W}$. Reliable negative labels and soft-negative labels are denoted by $Y^-$ and $Y^{\mathrm{soft}}$, respectively.

In the current implementation, masked BCE is the primary supervision term. To be consistent with the coarse loss formulation in the Methods, the detailed loss components are grouped into four categories: a supervised sparse-label loss $\mathcal{L}_{\text{sup}}$, a structural regularization loss $\mathcal{L}_{\text{struct}}$, an image-guided smoothness loss $\mathcal{L}_{\text{smooth}}$, and a regularization loss $\mathcal{L}_{\text{reg}}$. Specifically, the masked BCE losses for the area/interior and edge channels belong to $\mathcal{L}_{\text{sup}}$; the region-consistency and region-contrast losses belong to $\mathcal{L}_{\text{struct}}$; the image-guided smoothness term is $\mathcal{L}_{\text{smooth}}$; and the sparsity and parameter-regularization terms are grouped into $\mathcal{L}_{\text{reg}}$.

**Masked soft binary cross-entropy loss.** To accommodate sparse and partially reliable supervision, we use a masked soft binary cross-entropy loss. Positive labels are expanded using morphological dilation to define high-confidence regions, and each pixel $x$ is assigned a weight $W(x)$ that emphasizes annotated foreground, reliable background, and their immediate neighborhoods.

For a prediction channel $c \in \{A, E\}$, the masked BCE loss is written as

$$\mathcal{L}_{\text{bce}}^{c} = \sum_{x} W_c(x)\,\text{BCE}(\hat{Y}_c(x), Y_c(x)). \tag{26}$$

Here, $c = A$ denotes the area/interior channel and $c = E$ denotes the edge channel. The area channel is supervised by the sparse positive annotation mask, whereas the edge channel is supervised by the boundary mask derived from the same sparse annotation.

An additional foreground confidence term can be included to encourage confident activation within annotated foreground regions:

$$\mathcal{L}_{\text{push}}^{c} = -\frac{1}{|Y_c|} \sum_{x \in Y_c} P_c(x). \tag{27}$$

In practice, this term is included in the masked BCE implementation and is treated as part of the channel-wise masked classification loss.

**Region consistency loss.** To provide weak structural regularization, we compute a region consistency loss based on local image statistics. Let $\mu_I(x)$ and $\sigma_I^2(x)$ denote the local mean and variance of the reference image or feature map at pixel $x$, computed using a sliding window.

For a prediction channel $c$, a normalized soft foreground weight is defined as

$$w_c(x) = \frac{P_c(x)}{\sum_{x'} P_c(x') + \varepsilon}. \tag{28}$$

The variance consistency term is defined as

$$\mathcal{L}_{\text{var}}^{c} = \sum_{x} w_c(x) \left( \sigma_I^2(x) - \sum_{x'} w_c(x')\sigma_I^2(x') \right)^2, \tag{29}$$

and the mean consistency term is defined as

$$\mathcal{L}_{\text{mean}}^{c} = \sum_{x} w_c(x) \left( \mu_I(x) - \sum_{x'} w_c(x')\mu_I(x') \right)^2. \tag{30}$$

The channel-wise region consistency loss is

$$\mathcal{L}_{\text{cons}}^{c} = (1-\beta)\mathcal{L}_{\text{var}}^{c} + \beta\mathcal{L}_{\text{mean}}^{c}, \tag{31}$$

where $\beta$ balances the variance- and mean-based terms.

**Region contrast loss.** To encourage separation between predicted foreground and background regions, a region contrast loss is applied. For channel $c$, foreground and background statistics are computed as

$$\mu_f^c = \frac{\sum_x P_c(x)\mu_I(x)}{\sum_x P_c(x) + \varepsilon}, \qquad \mu_b^c = \frac{\sum_x (1 - P_c(x))\mu_I(x)}{\sum_x (1 - P_c(x)) + \varepsilon}, \tag{32}$$

$$(\sigma_f^c)^2 = \frac{\sum_x P_c(x)\sigma_I^2(x)}{\sum_x P_c(x) + \varepsilon}, \qquad (\sigma_b^c)^2 = \frac{\sum_x (1 - P_c(x))\sigma_I^2(x)}{\sum_x (1 - P_c(x)) + \varepsilon}. \tag{33}$$

The channel-wise contrast loss is defined as

$$\mathcal{L}_{\text{contrast}}^c = -\left[(1-\beta)\left\|(\sigma_f^c)^2 - (\sigma_b^c)^2\right\|_1 + \beta\left\|\mu_f^c - \mu_b^c\right\|_1\right]. \tag{34}$$

**Image-guided smoothness loss.** An image-guided smoothness loss is applied to the area/interior prediction to reduce fragmented activations while preserving image-defined boundaries. Edge-aware weights are defined as

$$g_x = \exp(-\gamma|\nabla_x I|), \qquad g_y = \exp(-\gamma|\nabla_y I|), \tag{35}$$

where $\gamma$ controls the sensitivity to image gradients.

The smoothness loss is given by

$$\mathcal{L}_{\text{smooth}} = \sum_x \left[g_x|\nabla_x P_A| + g_y|\nabla_y P_A|\right]. \tag{36}$$

This term is assigned a small weight in the default setting and acts as an auxiliary regularizer rather than a primary supervision signal.

**Sparsity and parameter regularization.** To suppress spurious predictions, a sparsity penalty is applied to the predicted probabilities:

$$\mathcal{L}_{\text{sparse}} = \frac{1}{2HW}\sum_x (P_A(x) + P_E(x)). \tag{37}$$

An $\ell_1$ regularization term is imposed on all network parameters $\theta$:

$$\mathcal{L}_{\ell_1} = \frac{1}{|\theta|}\sum_{p\in\theta}|p|. \tag{38}$$

The combined regularization term is

$$\mathcal{L}_{\text{reg}} = \mathcal{L}_{\ell_1} + \lambda_{\text{sparse}}\mathcal{L}_{\text{sparse}}. \tag{39}$$

**Overall objective.** The detailed loss components defined above correspond to the four coarse loss categories introduced in the Methods.

First, the supervised sparse-label loss is defined as

$$\mathcal{L}_{\text{sup}} = \alpha_A \mathcal{L}_{\text{bce}}^A + \alpha_E \mathcal{L}_{\text{bce}}^E, \tag{40}$$

where $A$ and $E$ denote the area/interior and edge channels, respectively. The area channel is supervised by the sparse positive annotation mask, whereas the edge channel is supervised by the boundary mask derived from the same sparse annotation. The optional foreground confidence term $\mathcal{L}_{\text{push}}^c$ is included in the channel-wise masked BCE implementation and is therefore treated as part of $\mathcal{L}_{\text{sup}}$.

Second, the structural regularization loss is defined as

$$\mathcal{L}_{\text{struct}} = \mathcal{L}_{\text{cons}} + \mathcal{L}_{\text{contrast}}, \tag{41}$$

where

$$\mathcal{L}_{\text{cons}} = \alpha_A \mathcal{L}^A_{\text{cons}} + \alpha_E \mathcal{L}^E_{\text{cons}}, \tag{42}$$

and

$$\mathcal{L}_{\text{contrast}} = \alpha_A \mathcal{L}^A_{\text{contrast}} + \alpha_E \mathcal{L}^E_{\text{contrast}}. \tag{43}$$

Thus, the region-consistency and region-contrast losses are grouped together as $\mathcal{L}_{\text{struct}}$, because both terms provide weak structural constraints based on local image or reference-map statistics.

Third, the image-guided smoothness term defined above is used directly as

$$\mathcal{L}_{\text{smooth}} = \sum_x \left[ g_x |\nabla_x P_A| + g_y |\nabla_y P_A| \right]. \tag{44}$$

This term acts on the area/interior prediction and encourages spatially coherent predictions while preserving image-defined boundaries.

Finally, the regularization loss is defined as

$$\mathcal{L}_{\text{reg}} = \mathcal{L}_{\ell_1} + \lambda_{\text{sparse}} \mathcal{L}_{\text{sparse}}, \tag{45}$$

where $\mathcal{L}_{\ell_1}$ penalizes model parameters and $\mathcal{L}_{\text{sparse}}$ penalizes excessive predicted activations.

The final training objective is therefore written as

$$\mathcal{L}_{\text{total}} = \lambda_{\text{sup}} \mathcal{L}_{\text{sup}} + \lambda_{\text{struct}} \mathcal{L}_{\text{struct}} + \lambda_{\text{smooth}} \mathcal{L}_{\text{smooth}} + \lambda_{\text{reg}} \mathcal{L}_{\text{reg}}. \tag{46}$$

In the reported experiments, the default weights were

$$\lambda_{\text{sup}} = 10, \quad \lambda_{\text{struct}} = 0.1, \quad \lambda_{\text{smooth}} = 0.1, \quad \lambda_{\text{reg}} = 0.05, \quad \lambda_{\text{sparse}} = 0.01. \tag{47}$$

Thus, the supervised masked BCE term dominates optimization, while the structural, smoothness, and regularization losses serve as weak auxiliary regularizers. The edge-channel BCE term contributes to $\mathcal{L}_{\text{sup}}$ when $\alpha_E > 0$.

Our ablation analysis shows that masked BCE is essential for learning meaningful target segmentation, whereas the auxiliary terms have weaker effects in the mitochondria benchmark. These additional terms are retained to support more challenging settings, such as low-contrast data, noisy sparse annotations, or weak boundary signals, but they can be reduced or disabled if they do not improve validation performance on a new dataset. This weighting strategy also helps maintain stable training by preventing auxiliary objectives from competing strongly with the primary sparse-supervision loss.

#### Shape- and geometry-based refinement of predicted labels

This section describes the post-processing and label refinement procedures applied to model predictions. The refinement pipeline combines shape similarity filtering for region interiors with geometric overlap constraints between boundary and area predictions. All operations are applied slice-wise in two dimensions unless otherwise stated.

##### Connected component extraction

Given a predicted binary mask volume $\hat{M} \in \{0,1\}^{D \times H \times W}$, connected components are extracted independently on each slice $z$ using 4-connectivity. Each connected component $C$ is characterized by a set of geometric and moment-based descriptors. Here, $M$ denotes the user-provided sparse positive annotation mask, whereas $\hat{M}$ denotes the predicted binary mask to be refined.

**Shape descriptor construction.** For each connected component $C$ in a two-dimensional slice, a shape descriptor vector $\mathbf{s}(C)$ is constructed by concatenating interpretable geometric features and low-order Hu moment invariants.

**Geometric features.** The geometric descriptor is defined as

$$\mathbf{g}(C) = \big[\text{circ}(C), \text{aspect}(C), \text{extent}(C), \text{solidity}(C), \text{eccentricity}(C), \text{Euler}(C)\big], \tag{48}$$

where circularity is computed as

$$\text{circ}(C) = \frac{4\pi A(C)}{P(C)^2 + \varepsilon}, \tag{49}$$

with $A(C)$ and $P(C)$ denoting the area and perimeter of the component, respectively. The aspect ratio is computed as the ratio between the major and minor axis lengths. Extent, solidity, eccentricity, and Euler number follow standard region-based definitions.

**Hu moment invariants.** Hu moment invariants are computed for each connected component. To reduce numerical instability from high-order moments in small sparse regions, we use only the first three Hu moments:

$$\mathbf{h}(C) = \big[h_1(C), h_2(C), h_3(C)\big]. \tag{50}$$

These Hu moments are not used as separate hard-thresholded features. Instead, they are included in the same descriptor vector and compared using relative deviation scores, as described below.

**Final shape vector.** The complete shape descriptor is given by

$$\mathbf{s}(C) = \big[\mathbf{g}(C) \parallel \mathbf{h}(C)\big]. \tag{51}$$

**Shape similarity filtering.** Let $\{\mathbf{s}_k^{\text{ref}}\}_{k=1}^{N_{\text{ref}}}$ denote the shape descriptors extracted from connected components in the sparse positive annotation mask $M$. For each predicted candidate component $C$, two relative deviation scores are computed.

First, we compute the template-wise deviation between the candidate component and the most similar annotated reference component:

$$E_{\text{template}}(C) = \min_k \max_i \frac{\left| s_i(C) - s_{k,i}^{\text{ref}} \right|}{\left| s_{k,i}^{\text{ref}} \right| + \varepsilon}. \tag{52}$$

This term evaluates whether the candidate component resembles at least one annotated reference component.

Second, we compute the deviation from the mean reference shape descriptor:

$$E_{\text{mean}}(C) = \max_i \frac{\left| s_i(C) - \mu_i^{\text{ref}} \right|}{|\mu_i^{\text{ref}}| + \varepsilon}, \tag{53}$$

where $\mu_i^{\text{ref}}$ is the mean value of the $i$-th descriptor dimension across all annotated reference components.

The two deviation values are converted into shape scores:

$$S_{\text{template}}(C) = -E_{\text{template}}(C), \qquad S_{\text{mean}}(C) = -E_{\text{mean}}(C). \tag{54}$$

Thus, scores closer to zero indicate higher similarity to the sparse annotated regions. The final shape score is defined as

$$S(C) = \max \left[ w_{\text{error}} S_{\text{template}}(C), w_{\text{range}} S_{\text{mean}}(C) \right], \tag{55}$$

where $w_{\text{error}}$ and $w_{\text{range}}$ control the relative contributions of template-wise similarity and mean-shape consistency. This formulation implements a permissive filtering criterion: a candidate component can obtain a high score if it is similar either to one annotated reference component or to the overall mean shape of the annotated regions.

Before ranking, simple topology and size constraints are applied to remove obvious false-positive regions. Candidate components with Euler number different from 1 are removed. In addition, the component area must satisfy

$$A_{\text{min}}^{\text{ref}} r_{\text{min}} \leq A(C) \leq \frac{A_{\text{max}}^{\text{ref}}}{r_{\text{min}}}, \tag{56}$$

where $A_{\text{min}}^{\text{ref}}$ and $A_{\text{max}}^{\text{ref}}$ are the minimum and maximum areas of the annotated reference components, and $r_{\text{min}}$ is the area tolerance ratio.

To focus the refinement step on newly discovered regions, candidate components that substantially overlap with existing sparse positive annotations are excluded. Specifically, a candidate component is skipped if more than 80% of its pixels already overlap with the annotation mask $M$ on the same slice.

The remaining candidate components are ranked in descending order of $S(C)$. Rather than applying a fixed absolute Hu-moment or shape-similarity threshold, SparseSeg adaptively determines the retained candidates from the score distribution within each dataset. Let $S_{\text{min}}$ and $S_{\text{max}}$ denote the minimum and maximum scores among all candidate components. The score cutoff is defined as

$$S_{\text{cutoff}} = S_{\text{max}} - (S_{\text{max}} - S_{\text{min}})\,\tau, \tag{57}$$

where $\tau$ is the shape-refinement threshold. Candidate components satisfying

$$S(C) \geq S_{\text{cutoff}} \tag{58}$$

are retained, subject to an upper bound on the maximum number of newly added regions. When $\tau \leq 0$, this shape-based filtering step is skipped. This adaptive selection strategy allows SparseSeg to expand sparse supervision in a dataset-specific manner while limiting excessive false-positive propagation.

##### Boundary–region consistency via bounding-box IoU

To enforce spatial consistency between predicted region interiors and boundary-like structures, a bounding-box overlap criterion is applied. For each region component $C_A$ and boundary component $C_L$ on the same slice, the intersection-over-union (IoU) of their bounding boxes is computed as

$$\text{IoU}(C_A, C_L) = \frac{|B(C_A) \cap B(C_L)|}{|B(C_A) \cup B(C_L)|}, \tag{59}$$

where $B(\cdot)$ denotes the axis-aligned bounding box.

Boundary components that exhibit excessive fill ratios within their own bounding boxes are discarded to suppress blob-like artifacts. A region component is retained if its maximum IoU with any valid boundary component exceeds a user-adjustable threshold. This threshold controls the strictness of boundary–region consistency filtering and can be tuned according to target morphology and annotation quality.

##### Volumetric consistency filtering

Following slice-wise refinement, retained components are optionally filtered in three dimensions by enforcing minimum and maximum volume constraints. Connected regions that are too thin along the axial direction or excessively large relative to reference annotations are removed.

##### Label augmentation

The final refined prediction volume is merged with existing annotations to form an expanded supervision set. These augmented labels are treated as reliable positives in subsequent training rounds, enabling iterative self-improvement while maintaining geometric consistency.

#### Benchmark settings

##### Evaluation metrics

We quantified segmentation performance using the raw Intersection-over-Union (IoU) and, for multi-model sparse-annotation benchmarks, the normalized relative IoU. Raw IoU was computed between the predicted binary mask and the corresponding expert annotation.

For benchmark settings in which the absolute IoU values varied substantially across models, datasets, or target structures, we additionally reported normalized relative IoU to facilitate comparison within the same benchmark. For each benchmark, all valid IoU values from all compared models were collected, and NaN values were excluded. Given a raw IoU value $I_{m,i}$ from model $m$ on sample $i$, the normalized relative IoU was computed as

$$I_{m,i}^{\mathrm{rel}} = \frac{\log(1 + 100 I_{m,i})}{\max_{m,i} \log(1 + 100 I_{m,i})}. \tag{60}$$

Here, the maximum was computed over all models and samples within the same benchmark setting. Thus, the best prediction in each benchmark was normalized to 1, while all other predictions were scaled relative to it.

##### Benchmark models and computing environment

We benchmarked SparseSeg under two complementary sparse-annotation settings. In the controlled label-sparsity benchmark, sparse labels were generated by randomly removing connected annotated regions or annotated slices at predefined sparsity ratios, including 30%, 50%, 70%, 80%, and 95%. This benchmark was repeated for 10 independent trials for each sparsity level to evaluate robustness to incomplete supervision.

In the ROI-level sparse-annotation benchmark, only a fixed number of positive ROIs, specifically 1, 5, or 10 ROIs, were retained as sparse target annotations. This more extreme sparse-label setting was repeated for five independent random ROI selections or initializations where applicable.

Across these benchmarks, we compared SparseSeg with several representative segmentation models, including organelle-specific EM segmentation models, general biomedical segmentation frameworks, dense volume-EM segmentation models, shape-prior-based instance segmentation methods, cryo-ET-oriented deep-learning tools, and an internal transformer-based variant of SparseSeg. The evaluated methods included MitoNet, nnU-Net, COSEM 2D/3D U-Net, DeePiCt, StarDist, and SparseSeg-ViT. Unless otherwise specified, the output of each model was binarized and evaluated using the same IoU-based evaluation pipeline.

All benchmark experiments, ablation tests, runtime measurements, and memory profiling were performed on the same workstation equipped with an AMD Ryzen Threadripper 7970X 32-core CPU operating at 4.00 GHz, 128 GB system RAM, an NVIDIA GeForce RTX 5080 GPU with 16 GB VRAM, and 5.46 TB local storage. To estimate the computational resource cost of SparseSeg, we recorded CPU utilization, process memory usage, system memory usage, GPU utilization, and GPU memory consumption during five representative SparseSeg trials. These measurements are summarized in Supplementary Tables 1 and 2. Because SparseSeg includes both neural-network training and non-network refinement steps, empirical wall-clock runtime and resource usage provide a practical estimate of computational efficiency.

**MitoNet.** MitoNet is a generalist deep-learning model developed for instance segmentation of mitochondria in electron microscopy images [6]. It was trained on diverse EM datasets and is designed to identify individual mitochondrial instances across different cellular contexts and imaging conditions. We included MitoNet as a strong organelle-specific benchmark for mitochondria segmentation.

For the MitoNet benchmark, we used the pretrained MitoNet_v1 model without task-specific sparse-label fine-tuning. Inference was performed using GPU acceleration with the quantized model enabled. The 2D inference parameters were set as follows: image downsampling = 1, segmentation confidence threshold = 0.20, center confidence threshold = 0.10, and minimum center distance = 6. For stack-level post-processing, the median filter size was set to 3, the minimum object size was set to 80 voxels, the minimum box extent was set to 2, and the maximum number of objects per class in 3D was set to 10,000. The inference plane was set to $yz$. Ortho-plane inference was disabled, label erosion and dilation were both set to 0, the voxel vote threshold was set to 1, and zarr storage was not used. Because this model was evaluated as a pretrained generalist mitochondria model, its result is reported as a reference baseline without sparse-label adaptation.

**nnU-Net.** nnU-Net is a self-configuring biomedical image segmentation framework based on the U-Net architecture [7]. For a given segmentation task, nnU-Net automatically configures key components of the pipeline, including preprocessing, network architecture, training strategy, and post-processing. We used nnU-Net as a strong general-purpose deep-learning baseline to evaluate whether the target-conditioned design of SparseSeg provides additional benefit beyond a well-optimized U-Net framework.

For the nnU-Net benchmark, sparse-label datasets were converted to the nnU-Net format and trained using the 3D full-resolution configuration. We used the ResEnc-M planner and plans, corresponding to `nnUNetPlannerResEncM` and `nnUNetResEncUNetMPlans`. Training was performed using all folds and a custom trainer with 300 epochs. Whole-volume prediction was then performed using the trained model, and only the final segmentation masks were saved for evaluation unless probability maps were explicitly required.

**COSEM 2D/3D U-Net.** We further compared SparseSeg with 2D and 3D U-Net baselines trained using the CellMap/COSEM-style segmentation pipeline. These models provide directly relevant dense EM segmentation baselines because they follow the standard U-Net design used for large-scale cellular EM segmentation.

For the 2D U-Net baseline, we used a 2D U-Net with input and target patch sizes of $80 \times 80$, batch size 8, learning rate $1 \times 10^{-4}$, 100 epochs, and 200 iterations per epoch. Training and prediction were performed for each benchmark sample using the corresponding sparse-label dataset split.

For the 3D U-Net baseline, we used a 3D U-Net with input and target patch sizes of $16 \times 80 \times 80$, batch size 2, learning rate $1 \times 10^{-4}$, 100 epochs, and 200 iterations per epoch. The 2D and 3D variants were included to evaluate the effect of slice-wise image context versus volumetric context under the same sparse-label benchmark setting.

**StarDist.** StarDist is an instance segmentation method that represents objects using star-convex polygons in 2D or star-convex polyhedra in 3D [8, 9]. It has been widely used for microscopy images containing densely packed cells or nuclei with approximately star-convex shapes. We included StarDist as a shape-prior-based baseline to test whether such geometric object representations are sufficient for segmenting cryo-vEM ultrastructures with more irregular morphologies.

For the ROI-level benchmark, StarDist was trained as a 3D instance segmentation model using the same raw volume and sparse labels. Binary sparse labels were first converted into instance labels by 3D connected-component labeling. We used 96 rays with anisotropy estimated from the training labels, a training patch size of $4 \times 80 \times 80$, batch size 16, 50 training epochs, learning rate $3 \times 10^{-4}$, and foreground-only sampling ratio 0.9. Whole-volume prediction was performed using tiled inference with $8 \times 8 \times 8$ tiles. Predictions were saved as instance-label TIFF volumes and then converted to binary masks for IoU evaluation.

**DeePiCt.** DeePiCt is a supervised deep-learning framework developed for mining molecular patterns and localizing macromolecular complexes in cellular cryo-electron tomography [10]. It combines 2D and 3D convolutional networks for segmentation of cellular compartments, continuous structures, and particle localization. Although DeePiCt was primarily designed for cryo-ET rather than cryo-vEM organelle segmentation, we included it as a recent cryo-EM-related deep-learning baseline to assess the transferability of cryo-ET-oriented segmentation frameworks to our target-conditioned cryo-vEM setting.

For DeePiCt, raw TIFF volumes and sparse-label TIFF masks were converted into MRC format before training and prediction. The ROI-level benchmark was run using 1, 5, and 10 positive ROIs with five repeated label initializations. For prediction, the 2D CNN configuration used a patch size of $288 \times 288$, crop size 48, normalization enabled, crop compensation enabled, and a prediction threshold of 0.5. Prediction patch dimensions were computed from the effective patch size after crop compensation. Each benchmark sample was trained with a separate configuration file and model output path to avoid checkpoint reuse across runs. The resulting MRC predictions were converted back to binary TIFF volumes for evaluation.

**SparseSeg-ViT.** To evaluate whether the performance of SparseSeg depends specifically on the multi-kernel U-Net backbone, we constructed an internal architectural baseline termed SparseSeg-ViT. SparseSeg-ViT follows the same target-conditioned training strategy, sparse supervision setting, loss functions, and refinement pipeline as SparseSeg, but replaces the U-Net backbone with a Vision Transformer-based backbone [11]. This comparison isolates the contribution of the convolutional multi-kernel design from the remaining components of the SparseSeg framework. SparseSeg-ViT was trained and evaluated using the same ROI-level sparse-label settings, refinement procedure, and evaluation metrics as SparseSeg.

**Computing environment.** All benchmark experiments, ablation tests, runtime measurements, and memory profiling were performed on the same workstation equipped with an AMD Ryzen Threadripper 7970X 32-core CPU operating at 4.00 GHz, 128 GB system RAM, an NVIDIA GeForce RTX 5080 GPU with 16 GB VRAM, and 5.46 TB local storage. Unless otherwise specified, all reported training time, inference time, and peak memory usage were measured on this machine. Runtime varied depending on the benchmark model, volume size, patch size, and number of sparse-label repeats. For fair comparison, all models were evaluated on the same benchmark volumes and sparse-label settings wherever applicable.

#### Construction of Datasets with Controlled Label Sparsity

To systematically evaluate model performance under varying degrees of annotation sparsity, we constructed a series of sparsified 3D datasets from fully annotated volumetric labels. The sparsification procedure was designed to preserve local morphological coherence while simulating realistic incomplete annotation scenarios commonly encountered in large-scale volumetric imaging.

**Source data and biological scope.** All volumetric datasets used in this study were derived from the Janelia COSEM project, which provides high-resolution, fully annotated 3D electron microscopy volumes of whole cells. We selected three representative cell types—*HeLa* cells (hela2), *Jurkat* T lymphocytes (jurkat), and macrophages (macrophage)—covering diverse cellular morphologies and ultrastructural organizations. For each cell type, we considered six major organelle classes, including mitochondria (mito), endoplasmic reticulum (er), endosomes (endo), lysosomes (lyso), Golgi apparatus (golgi), and the cell nucleus (nucleus). This combination of cell types and organelles enables systematic evaluation of model robustness and generalization across both cellular and subcellular domains.

**Uniform intensity-based partitioning and connected-component labeling.** Given a 3D annotation volume $V \in \mathbb{R}^{D\times H\times W}$, voxel intensities were first partitioned into $N$ uniformly spaced intervals along the global intensity range, with $N = 8$ used in all experiments. For each interval, a binary mask was generated by thresholding voxels whose values fell within the corresponding intensity bounds. Three-dimensional connected-component labeling (26-connectivity) was then applied independently within each interval. To ensure global uniqueness, labels obtained from different intervals were assigned non-overlapping identifiers. This procedure resulted in a set of spatially coherent 3D regions spanning the entire intensity distribution of the original annotation volume.

**Region-level sparsification by random removal.** To generate datasets with controlled levels of sparsity, we introduced region-level label removal. For volumes containing a large number of connected regions (typically more than 500 regions), sparsity was implemented by randomly selecting a fraction of region labels and removing them entirely. Specifically, for a target sparsity ratio $r \in \{30, 50, 70, 80, 95\}\%$, a corresponding fraction $r/100$ of region labels was sampled uniformly without replacement and reassigned to background (label 0). This strategy mimics incomplete annotation scenarios in which entire structures are absent from the labeled data.

For the extreme ROI-level benchmark, we used a similar sparsification pipeline but retained a fixed number of annotated ROIs, specifically 1, 5, or 10 ROIs, instead of retaining labels according to a sparsity ratio.

**Slice-wise sparsification for low region counts.** For volumes with a smaller number of connected regions, removing entire regions would lead to excessive information loss. In these cases, a slice-wise sparsification strategy was adopted. For each connected region, a subset of its occupied axial ($z$) slices was randomly selected according to the target sparsity ratio, and voxels belonging to the region were removed only within those slices. The remaining slices of the same region were preserved. This approach maintains partial structural continuity and better reflects realistic slice-dependent annotation gaps observed in serial imaging workflows.

**Binary mask generation.** After sparsification, all non-zero labels were converted into binary masks by assigning foreground voxels a value of 1 and background voxels a value of 0. The resulting binary masks were saved independently for each sparsity level and used as sparse supervision signals for downstream training and evaluation.

**Summary.** Using this procedure, we generated a family of datasets with systematically controlled annotation sparsity across multiple cell types and organelles. By varying the sparsity ratio while preserving region morphology and spatial coherence, this framework enables rigorous evaluation of model robustness and generalization under progressively reduced supervision.

#### Benchmarking across organelles and cell types

We further benchmarked SparseSeg against StarDist across multiple cell types, including Jurkat T cells, HeLa cells (hela2), and macrophages, and across a diverse set of subcellular targets, including mitochondria, endoplasmic reticulum (ER), Golgi apparatus, lysosomes, nucleus, and endosomes. Overall, SparseSeg outperformed StarDist in the majority of cases, demonstrating consistently higher robustness under varying levels of label sparsity and across heterogeneous cellular contexts.

We quantify segmentation performance using the intersection over union (IoU) and the false positive rate (FPR), defined as

$$\mathrm{IoU} = \frac{|G \cap P|}{|G \cup P|}, \quad \mathrm{FPR} = \frac{\sum(\neg G \wedge P)}{\sum(\neg G \wedge P) + \sum(\neg G \wedge \neg P)}, \tag{61}$$

where $G$ and $P$ denote the ground-truth and predicted binary masks, respectively.

However, for a small subset of organelle–cell-type combinations—notably lysosomes in Jurkat cells, lysosomes and ER in HeLa cells, and ER and Golgi apparatus in macrophages—StarDist achieved comparable performance and, in some masking regimes, marginally outperformed SparseSeg. Notably, these cases correspond to organelles for which the reference annotations provided by the Janelia COSEM project exhibit substantially higher noise levels, including fragmented labels, ambiguous boundaries, and inconsistent coverage across slices. In these settings, both models perform noticeably worse than in mitochondrial benchmarks, where annotations are generally of higher quality and consistency.

These observations suggest that the apparent performance parity in these cases primarily reflects limitations in annotation quality rather than intrinsic advantages of the competing method. Indeed, both SparseSeg and StarDist struggle to recover biologically coherent segmentations when supervision is dominated by label noise, underscoring the sensitivity of quantitative benchmarking to annotation fidelity.

Since ilastik is not designed for high-throughput or batch processing of large volumetric datasets, its practical applicability for systematic benchmarking across multiple organelles, cell types, and sparsity conditions is inherently limited. This motivated its exclusion from large-scale comparative analyses.

Additional benchmarking results and representative comparisons are summarized in Supplementary Fig. 2, which further illustrate the relative strengths and failure modes of SparseSeg and StarDist across challenging segmentation scenarios.

#### Hyperparameter sensitivity and ablation analysis

We performed ablation and robustness analyses to evaluate the contribution and sensitivity of several key design choices in SparseSeg. These analyses included the shape-refinement threshold, kernel configuration, boundary–region IoU threshold, low-weight coefficient, patch size, U-Net backbone structure, loss-component combinations, and feature-group ablations. For the feature ablation, the extracted feature channels were grouped into three major categories: structural filters, frequency filters, and Wiener-denoised features. All hyperparameter sensitivity and ablation experiments in this section were performed on the HeLa mitochondria benchmark. Unless explicitly varied in each experiment, all other parameters were kept at the default settings used in the SparseSeg pipeline, including `roi num` =10, `patch_scale`=80, `area_coef`=1.0, `edge_coef`=1.0, `iou_thresh`=0.6, `threshold`=0.01, `negative_threshold`=3, `low_weight_coeff`=50, and `sparsity_weight`=1.0.

As shown in Supplementary Fig. 3a, the shape-refinement threshold strongly affected segmentation performance. Compared with the control setting without shape-based refinement, nonzero refinement thresholds generally improved relative IoU, indicating that geometry-based candidate selection helps suppress unreliable predictions and improve label propagation. The best-performing threshold varied across trials, reflecting the dependence of this parameter on target morphology, image contrast, and annotation quality.

We next evaluated the effect of kernel size and the number of kernels (Supplementary Fig. 3b). Different kernel configurations all achieved competitive performance, but the optimal setting varied across trials. Single-kernel and multi-kernel configurations captured different contextual scales, and using more kernels did not always guarantee the highest relative IoU for every individual dataset. We therefore revised the interpretation of the multi-kernel design to avoid overstating its advantage. The multi-kernel architecture is intended to provide a general mechanism for integrating information across multiple spatial scales, which may be useful for targets that differ in size, boundary sharpness, and morphological complexity across vEM datasets, rather than being guaranteed to improve performance for every single target structure.

The boundary–region IoU threshold also affected performance (Supplementary Fig. 3c). Moderate thresholds generally maintained high performance, whereas the control setting with an IoU threshold of 1.0 showed reduced performance and increased variability. These results support the contribution of boundary–region consistency filtering while also indicating that the strictness of refinement should be tuned according to the dataset. In practice, overly strict filtering may remove true target regions with atypical morphology, whereas overly permissive filtering may allow false-positive pseudo-labels to propagate.

The low-weight coefficient influenced segmentation performance, but its effect was more variable across settings (Supplementary Fig. 3d). This coefficient controls the down-weighting of regions far from positive annotations. Therefore, its optimal value depends on the spatial distribution of user-provided labels, target density, and the degree of background heterogeneity. In the tested mitochondria benchmark, intermediate-to-large values generally produced competitive performance, but no single value was uniformly optimal across all trials.

Patch size was another important user-configurable parameter. As shown in Supplementary Fig. 3e, a very small patch size, such as `patch_scale`=20, produced the lowest relative IoU, indicating that insufficient spatial context limits the ability of the model to distinguish the target structure from surrounding ultrastructure. Increasing the patch size generally improved performance by providing more contextual information. However, larger patches also showed higher variability, likely because they include more heterogeneous background and neighboring structures. These results show that SparseSeg is sensitive to patch size and that this parameter should be selected according to the expected target size and local context. In practice, we recommend choosing a patch size comparable to or moderately larger than the characteristic size of the target structure, and then adjusting it according to memory constraints and validation performance.

We also evaluated the effect of the U-Net backbone structure (Supplementary Fig. 3f). Ablating U-Net layers reduced segmentation performance, indicating that sufficient encoder–decoder depth and hierarchical feature aggregation are important for robust sparse-label segmentation. This result supports the use of the U-Net backbone in SparseSeg, because the model needs to combine local boundary cues with broader contextual information to distinguish target structures from surrounding ultrastructure.

We further performed loss-component ablation experiments (Supplementary Fig. 3g). Removing the supervised masked BCE loss caused the network to fail to learn meaningful target segmentation, confirming that masked BCE is the essential supervised loss term under sparse annotation. In contrast, removing the structural regularization, smoothness, or parameter regularization terms did not substantially reduce performance in this mitochondria benchmark, and in some cases slightly improved relative IoU. This result is consistent with the fact that these auxiliary terms are assigned much smaller weights than the supervised BCE loss in the default configuration. Specifically, the default loss weights are $\lambda_{\text{sup}} = 10$, $\lambda_{\text{struct}} = 0.1$, $\lambda_{\text{smooth}} = 0.1$, and $\lambda_{\text{reg}} = 0.05$. Therefore, these terms should be interpreted as weak regularizers rather than primary supervision signals. They are retained in SparseSeg because they may provide additional constraints in more challenging settings, such as low-contrast images, noisy sparse annotations, or targets with weak boundaries. However, for relatively stable targets, users may reduce or disable these auxiliary weights if validation performance does not improve.

Finally, we evaluated the contribution of different feature groups (Supplementary Fig. 3h). Removing any of the three feature groups led to a clear decrease in relative IoU compared with the full-feature control. Ablating structural filtering responses reduced the model's ability to capture multi-scale morphological and boundary-related cues. Removing frequency-domain features also decreased performance, indicating that global and band-limited image information provides complementary context for target identification. Ablating Wiener-denoised features caused a similar performance drop, suggesting that denoising-based representations help stabilize learning under low-contrast and noisy vEM imaging conditions. These results indicate that the three handcrafted feature groups are not redundant but instead contain complementary information that collectively improves the representational capacity of the network.

Together, these analyses show that SparseSeg performance depends on both input feature representation and user-configurable model parameters. Feature-group ablations demonstrate that structural filters, frequency filters, and Wiener-denoised representations provide complementary information for segmentation. Among user-configurable parameters, SparseSeg is most sensitive to parameters controlling spatial context and refinement strictness, especially `patch_scale`, the shape-refinement threshold, and the boundary–region IoU threshold. In contrast, several auxiliary regularization terms have weaker effects under the tested mitochondria benchmark. Based on these results, we recommend that users first use the full feature representation, then tune `patch_scale` according to the target size and available memory, and finally adjust the refinement thresholds to balance false-positive suppression and recall. Auxiliary loss weights should be kept small by default and only increased when dataset-specific validation suggests that additional structural or smoothness constraints improve performance.

##### Limitations and future improvements

SparseSeg is designed as a target-conditioned segmentation framework, and therefore its predictions can be influenced by the positive annotations provided by the user. As shown in Supplementary Fig. 4, different sets of positive labels on the same raw image can lead to different segmentation results. In trial 3, the model identifies more candidate target structures, but this also introduces more false-positive predictions compared with trial 1. This variability is likely caused by the heterogeneity of the input positive labels. When the annotated regions cover only a limited subset of the target morphology, the model may learn a biased representation of the target structure and propagate this bias during iterative refinement.

This limitation reflects an inherent challenge of sparse target-conditioned segmentation: the quality and representativeness of the initial positive annotations directly affect the reliability of the expanded supervision. Although the refinement and filtering steps in SparseSeg reduce obvious false positives, the current framework cannot fully eliminate errors caused by incomplete or non-representative input labels. In future work, this limitation may be addressed by incorporating foundation segmentation models or promptable vision models, such as SAM-based architectures [12–14], to provide stronger pretrained image priors and improve robustness to sparse or heterogeneous annotations. Such models could potentially serve as initialization, auxiliary pseudo-label generators, or interactive refinement modules within the SparseSeg pipeline.

#### Implementation Pipeline and Usage Guide

This Supplementary provides practical implementation details for applying the proposed segmentation framework, including input preparation, optional preprocessing, and training parameter configuration. These details are intended to facilitate reproducibility and user adoption, and are complementary to the methodological description presented in the main text.

##### Input data preparation and optional preprocessing

The segmentation framework requires two mandatory volumetric inputs and one optional input:

- **Input image volume**: the volumetric EM dataset to be segmented.

- **Sparse label mask**: a binary volume indicating sparsely annotated target structures.
- **Background mask (optional)**: a binary volume indicating regions known *not* to contain the target structure.

All input volumes must share identical spatial dimensions along the $x$, $y$, and $z$ axes and be stored in `TIFF` stack format. Input files should be placed in the `inputdata` directory. If a background mask is provided, it must follow the naming convention `negative_<filename>`, where `<filename>` corresponds to the raw image or label mask.

Input volumes can be generated using the provided Fiji/ImageJ macro (`preprocessing.ijm`), which performs stack alignment, canvas normalization, and ROI-based label extraction. Alternatively, users may prepare these volumes using custom pipelines, provided that the resulting data conform to the required format and dimensional consistency.

##### Model training and parameter configuration

Once the required input volumes are prepared, model training can be initiated by running the script `iterative_seg.py`. The training process supports extensive parameter customization to accommodate diverse organelle morphologies, image characteristics, and annotation sparsity levels.

Key configurable parameters include:

- `z_threshold`: maximum number of consecutive $z$ slices that a target structure is allowed to occupy. This parameter helps suppress elongated intercellular or non-target structures that span excessive axial depth.
- `iou_thresh`: minimum required overlap ratio between predicted edge regions and corresponding interior regions. Larger values enforce sharper, well-defined boundaries.
- `threshold`: morphological similarity threshold between predicted regions and provided sparse labels, computed using 2D shape descriptors. Higher values restrict predictions to regions more closely matching annotated morphology.
- `area_coef` and `edge_coef`: weighting coefficients controlling the relative importance of area-based versus edge-based cues during training. Targets with strong membrane definition benefit from higher `edge_coef` values.
- `negative_threshold`: fraction of sampled training patches drawn from negative or background regions. Increasing this parameter encourages sparser predictions and reduces false positives.
- `low_weight_coeff`: spatial clustering constraint reflecting the expected physical size of target structures. For compact targets forming clusters of approximately $10 \times 10 \times 10$ voxels, setting `low_weight_coeff` to 10 suppresses isolated noise. For diffuse or weakly clustered targets, larger values are recommended to preserve distant signals.
- `sparsity_weight`: explicit sparsity regularization coefficient applied to prediction probabilities. Larger values promote sparser segmentation outputs.
- `patch_scale`: characteristic spatial scale of extracted training patches and the most critical parameter for successful segmentation. This value should be comparable to or slightly larger than the expected target size. For example, for a target with approximate dimensions of $30 \times 50 \times 30$ voxels, a `patch_scale` in the range of 50–100 is recommended.

##### Iterative training and label refinement

The framework supports iterative training through the parameter `params_list`, which specifies a sequence of training iterations. For example:

```
params_list = [
    {"iteration_idx": 0},
    {"iteration_idx": 1},
    {"iteration_idx": 2}
]
```

In this configuration, training is performed for three successive iterations. After each iteration, refined predictions from the previous round are incorporated as updated supervision for the next training stage. Users may optionally assign different parameter values to each iteration to progressively adjust sparsity constraints, patch scale, or morphological thresholds.

In most practical applications, satisfactory performance can be achieved by tuning only `sparsity_weight` and `patch_scale`, while keeping other parameters fixed. The iterative refinement strategy enables robust segmentation from extremely sparse annotations and improves convergence stability across diverse datasets.

### Supplementary References

[11] Alexey Dosovitskiy, Lucas Beyer, Alexander Kolesnikov, Dirk Weissenborn, Xiaohua Zhai, Thomas Unterthiner, Mostafa Dehghani, Matthias Minderer, Georg Heigold, Sylvain Gelly, Jakob Uszkoreit, and Neil Houlsby. An image is worth 16x16 words: Transformers for image recognition at scale. In *International Conference on Learning Representations*, 2021.

[12] Alexander Kirillov, Eric Mintun, Nikhila Ravi, Hanzi Mao, Chloe Rolland, Laura Gustafson, Tete Xiao, Spencer Whitehead, Alexander C. Berg, Wan-Yen Lo, Piotr Dollár, and Ross Girshick. Segment anything. In *Proceedings of the IEEE/CVF International Conference on Computer Vision*, pages 4015–4026, 2023.

[13] Jun Ma, Yuting He, Feifei Li, Lin Han, Chenyu You, and Bo Wang. Segment anything in medical images. *Nature Communications*, 15:654, 2024. doi: 10.1038/s41467-024-44824-z.

[14] Nikhila Ravi, Valentin Gabeur, Yuan-Ting Hu, Ronghang Hu, Chaitanya Ryali, Tengyu Ma, Haitham Khedr, Roman Rädle, Chloe Rolland, Laura Gustafson, Eric Mintun, Junting Pan, Kalyan Vasudev Alwala, Nicolas Carion, Chao-Yuan Wu, Ross Girshick, Piotr Dollár, and Christoph Feichtenhofer. SAM 2: Segment anything in images and videos. *arXiv preprint arXiv:2408.00714*, 2024.
